# Protein tyrosine phosphatase receptor kappa regulates glycolysis and *de novo* lipogenesis to promote hepatocyte metabolic reprogramming in obesity

**DOI:** 10.1101/2023.12.01.569004

**Authors:** Eduardo H. Gilglioni, Ao Li, Wadsen St-Pierre-Wijckmans, Tzu-Keng Shen, Israel Pérez-Chávez, Garnik Hovhannisyan, Michela Lisjak, Javier Negueruela, Valerie Vandenbempt, Julia Bauzá-Martinez, Jose M. Herranz, Daria Ezerina, Stephane Demine, Zheng Feng, Thibaut Vignane, Lukas Otero-Sánchez, Flavia Lambertucci, Alena Prašnická, Jacques Devière, David C. Hay, Jose A. Encinar, Sumeet Pal Singh, Joris Messens, Milos R. Filipovic, Hayley J. Sharpe, Eric Trépo, Wei Wu, Esteban N. Gurzov

## Abstract

Fat accumulation, *de novo* lipogenesis, and glycolysis are key drivers of hepatocyte reprogramming and the consequent metabolic dysfunction-associated steatotic liver disease (MASLD). Here we report that obesity leads to dysregulated expression of hepatic protein-tyrosine phosphatases (PTPs). PTPRK was found to be increased in steatotic hepatocytes in both humans and mice, and positively correlated with PPARγ-induced lipogenic signalling. High-fat-fed PTPRK knockout mice displayed reduced weight gain and hepatic fat accumulation. Phosphoproteomic analysis in primary hepatocytes and hepatic metabolomics identified fructose-1,6-bisphosphatase 1 and glycolysis as PTPRK targets in metabolic reprogramming. Silencing PTPRK in hepatoma cell lines resulted in reduced colony-forming ability and PTPRK knockout mice developed smaller tumours after diethylnitrosamine-induced hepatocarcinogenesis. Our study defines a novel role for PTPRK in regulating hepatic glycolysis, lipid metabolism, and tumour development. PTPRK inhibition may provide therapeutic possibilities in obesity-associated liver diseases.

**Highlights:**

- Hepatic receptor-type PTPs are increased in MASLD
- PTPRK is expressed in hepatocytes and upregulated in obesity
- PTPRK deficiency reduces body fat mass and liver steatosis in diet-induced obesity
- PTPRK regulates hepatic glycolysis and lipogenesis, promoting tumorigenesis

## Introduction

Consumption of processed industrialized foods with high caloric density and reduced energy expenditure results in nutrient overload. In response, cells adapt by storing energy in the form of triglycerides, generating adipose tissue expansion and ectopic fat deposition in organs such as the liver. The presence of a lipid-rich environment has deleterious pathological consequences, including insulin resistance and dyslipidaemia. If not resolved, it can evolve into non-alcoholic fatty liver disease (NAFLD), recently renamed metabolic dysfunction-associated steatotic liver disease (MASLD), affecting a quarter of the global adult population^1,2^. Non-alcoholic steatohepatitis (NASH), a severe necro-inflammatory form of NAFLD/MASLD, poses a significant health problem^3,4^. Moreover, NAFLD/MASLD has emerged as a leading cause of hepatocellular carcinoma (HCC), a highly heterogeneous and aggressive malignancy^5^.

The liver is central to nutrient sensing and has a significant impact during obesity, resulting in abnormal lipid accumulation and hepatocyte metabolic reprogramming. This involves the intricate reorganization of anabolic and catabolic processes, all under the transcriptional control of nutrient-sensitive receptors. In the context of obesity and HCC, several transcription factors, including PPARs, SREBP1c, ChREBP, and HIF, participate in the reshaping of metabolic pathways^3,4^. In addition, post-translational modifications, including phosphorylation of protein tyrosine residues, are dysregulated in nutrient overload and obesity^6^.

Hepatic expression of Protein Tyrosine Phosphatases (PTPs) is affected in steatotic livers and NASH. PTPs were conventionally perceived as enzymes responsible for terminating or modulating signals initiated by tyrosine kinases^7^. Accumulating evidence reveals their potential as signal propagators^6,8^. For example, PTPN2 facilitates signalling through both STAT1 and STAT3, exerting distinct influences on NASH and HCC^9^. Oxidative inactivation of PTPN2 induces an insulin-STAT5-IGF-1-growth hormone pathway in conditions of selective insulin resistance, contributing to obesity^10^. Receptor-type PTPs (RPTPs) are transmembrane enzymes adapted to sense and transduce extracellular cues into intracellular catalytic events^11^. In obesity, inflammatory signals induce the expression of PTPRG in the liver^12^. Deleting or overexpressing PTPRG enhances or suppresses hepatic insulin sensitivity, respectively^12^. However, a lack of comprehensive studies to understand liver PTPomes in obesity means that the role of PTPs in regulatory mechanisms and their potential use as biomarkers or therapeutic targets remain largely unexplored.

PTPRK is a transmembrane receptor belonging to the R2B subfamily of RPTPs, known to engage in homophilic interactions and localises to cell-cell contacts^11^. Cell adhesion proteins have been proposed as PTPRK substrates, and accumulating data associate PTPRK with several human diseases^13-15^. The regulation of PTPRK involves a proteolytic cascade (furin, ADAM10, and γ-secretase), potentially releasing the intracellular catalytic domain to interact and dephosphorylate proteins in the cytoplasm or nucleus^16^. PTPRK is transcriptionally regulated by transforming growth factor-β (TGF-β) and Notch signalling^17^. Despite its potential in cell signalling, the downstream events regulated by PTPRK remain unknown. In this study, we demonstrated that PTPRK is upregulated in fatty hepatocytes and investigated its role in obesity-associated liver dysfunction. PTPRK deficiency leads to severe metabolic changes in hepatocytes, culminating in reduced diet-induced obesity and hepatic fat accumulation in mice. We identified fructose-1,6-bisphosphatase 1 (FBP1), a gluconeogenic enzyme, as PTPRK target, bearing significant implications for glucose metabolism and liver tumour growth. These findings underscore the pivotal role of PTPRK as key driver in the metabolic reprogramming of hepatocytes induced by obesity.

## Results

### Hepatic PTP expression is dysregulated in obesity-associated liver dysfunction

The presence and potential contribution of PTP expression in the progression to NAFLD/MASLD in obesity and the development of HCC remains unknown. Therefore, we conducted a comprehensive analysis of the complete proteome and PTP expression patterns, using liquid chromatography-tandem mass spectrometry (LC-MS/MS) analysis of human liver samples (**Fig. 1A**). The cohort included liver biopsies obtained from individuals exhibiting varying degrees of liver disease, encompassing simple steatosis (Metabolic Associated Fatty Liver (MAFL)), NASH, HCC, and control samples from individuals without evidence of liver damage (healthy liver). The heatmap illustrating the total proteome alterations showed variations in protein expression and sample heterogeneity across different stages of liver dysfunction (**Supplementary Fig. 1A**). KEGG pathway analyses revealed that several protein modifications are related to metabolic dynamics. When comparing steatosis with healthy livers, we observed activation of oxidative phosphorylation, starch, and sucrose metabolism. We also observed activation of glutathione metabolism, alongside the suppression of the tight junction pathway (**Supplementary Fig. 1B**). Activated pathways in NASH included ECM receptor interaction, oxidative phosphorylation, and focal adhesion (**Supplementary Fig. 1C**). Conversely, pathways related to the pentose phosphate pathway (PPP), purine metabolism, histidine metabolism, and cysteine and methionine metabolism were suppressed. Further, comparing NASH to steatosis, oxidative phosphorylation was suppressed in NASH, while ECM receptor interaction, focal adhesion, and ribosome pathways were activated (**Supplementary Fig. 1D**). Proteome analysis revealed key enzymes involved in fatty acid uptake and metabolism, CD36, CPT1, and SCD, upregulated in steatosis and NASH samples (**Fig. 1B**). Among the samples, 18 out of the 37 human PTP proteins were identified (**Fig. 1C**). Analysis of the PTPome revealed that samples within the same disease stage displayed similar PTP expression patterns (**Fig. 1D**), while analysis of RPTPs and non-receptor protein tyrosine phosphatases (PTPNs) revealed opposing patterns across the stages of the disease (**Fig. 1E**). Several PTPNs were downregulated with disease, while RPTPs generally exhibited low expression levels in healthy liver samples but showed a marked upregulation in steatosis and NASH. Specifically, PTPRK, PTPRE, PTPRM, PTPRF, and PTPRA were elevated in steatosis and NASH (**Fig. 1F**). Single-cell RNA sequencing of healthy-obese livers^18^ revealed that PTPRK is the most abundant RPTP in hepatocytes, followed by PTPRG and PTPRM, with PTPRE mainly found in dendritic cells (**Fig. 1G, Supplementary Fig. 1E**). Additionally, RPTP displayed comparable mRNA patterns in the E-MEXP-3291 dataset (**Fig. 1H**). Correlation analysis showed that hepatic PTPRK, PTPRG, and PTPRE transcript levels positively correlate with PPARγ (**Fig. 1I**), a master regulator of lipid accumulation in hepatocytes. Immunohistochemistry (IHC) analyses in human liver samples showed that PTPRK levels were higher in steatosis and NASH, whereas healthy liver displayed relatively lower expression. The PTPRK intracellular domain localised within various cellular regions, including the nucleus of steatotic hepatocytes (**Fig. 1J**). The striking remodelling of PTPomes with disease onset indicates a potential causative role in fat accumulation and liver dysfunction.

**Figure 1.**
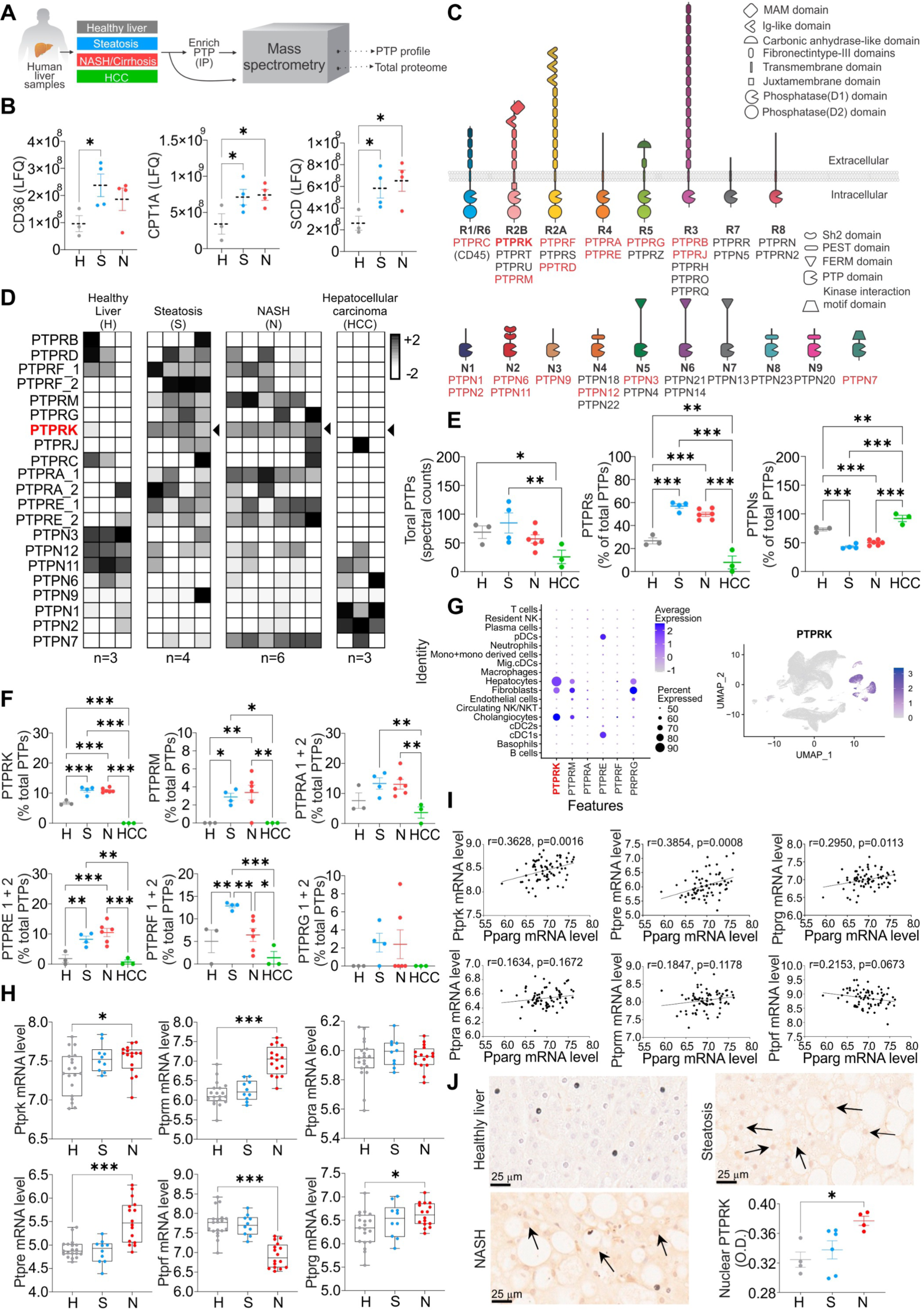
Enhanced PTPRK expression in human livers with steatosis and non-alcoholic steatohepatitis (NASH). (**A**) Methodological approach schematic illustrating the quantification of protein tyrosine phosphatase (PTP) profile and total proteome in human livers. (**B**) Quantification of lipid metabolism-related proteins using label-free quantification (LFQ) in healthy (H), steatotic (S) and NASH (N) livers. (**C**) Schematic representation of PTP families and their characteristic domains. PTPs detected by mass spectrometry are labelled in red. (**D**) Heat map displaying the hepatic PTP profile. (**E**) Spectral counts of total PTPs and the proportional contribution of receptor and non-receptor PTPs among the identified PTPs. (**F**) The proportional contribution of PTPRK and other receptor PTPs to the total identified PTPs is shown. (**G**) Data extracted from GSE192740 showing hepatocyte expression of PTPRK. (**H**) RPTP mRNA levels in the E-MEXP-3291 dataset. (**I**) Correlation analysis between RPTPs and Pparg mRNA levels in the E-GEOD-48452 dataset. (**J**) Representative immunohistochemistry (IHC) images displaying PTPRK staining and quantitative results for nuclear PTPRK. Statistical significance is denoted as \**p*<0.05, \*\**p*<0.01, \*\*\**p*<0.001.

### Hepatocyte PTPRK is induced in obesity and positively correlates with PPARγ in mouse models and primary hepatocytes

To gain functional insights into the hepatic role of PTPRK we took advantage of obesogenic mouse models. C57BL/6N mice were exposed to diet-induced obesity over a 12- week period. Both high-fat diet (HFD) and high-fat, high-fructose high-cholesterol diet (HFHFHCD) resulted in a noticeable increase in body weight (**Fig. 2A**), primarily attributed to increased body fat mass (**Fig. 2B**). This was accompanied by elevated fasting insulin levels (**Fig. 2C**), glucose intolerance (**Fig. 2D**), and reduced insulin sensitivity (**Fig. 2E**). PTPRK is expressed in hepatocytes, but undetectable in subcutaneous and visceral adipose tissues or muscle (**Fig. 2F**). Mice fed with HFHFHCD exhibited a greater liver weight, liver-to-body weight ratio, and liver fat mass compared to the control group (**Fig. 2G,H**), consistent with a more advanced stage of fatty liver development. PTPRK protein expression is enhanced in HFD and HFHFHCD-fed mice livers, accompanied by PPARγ upregulation (**Fig. 2I**). In line with these results, adenovirus-mediated overexpression of PTPRK in mouse livers resulted in a concomitant increase in PPARγ levels (**Fig. 2J**). These results demonstrate a consistent involvement of PTPRK in lipid metabolism and diet-induced liver dysfunction.

**Figure 2.**
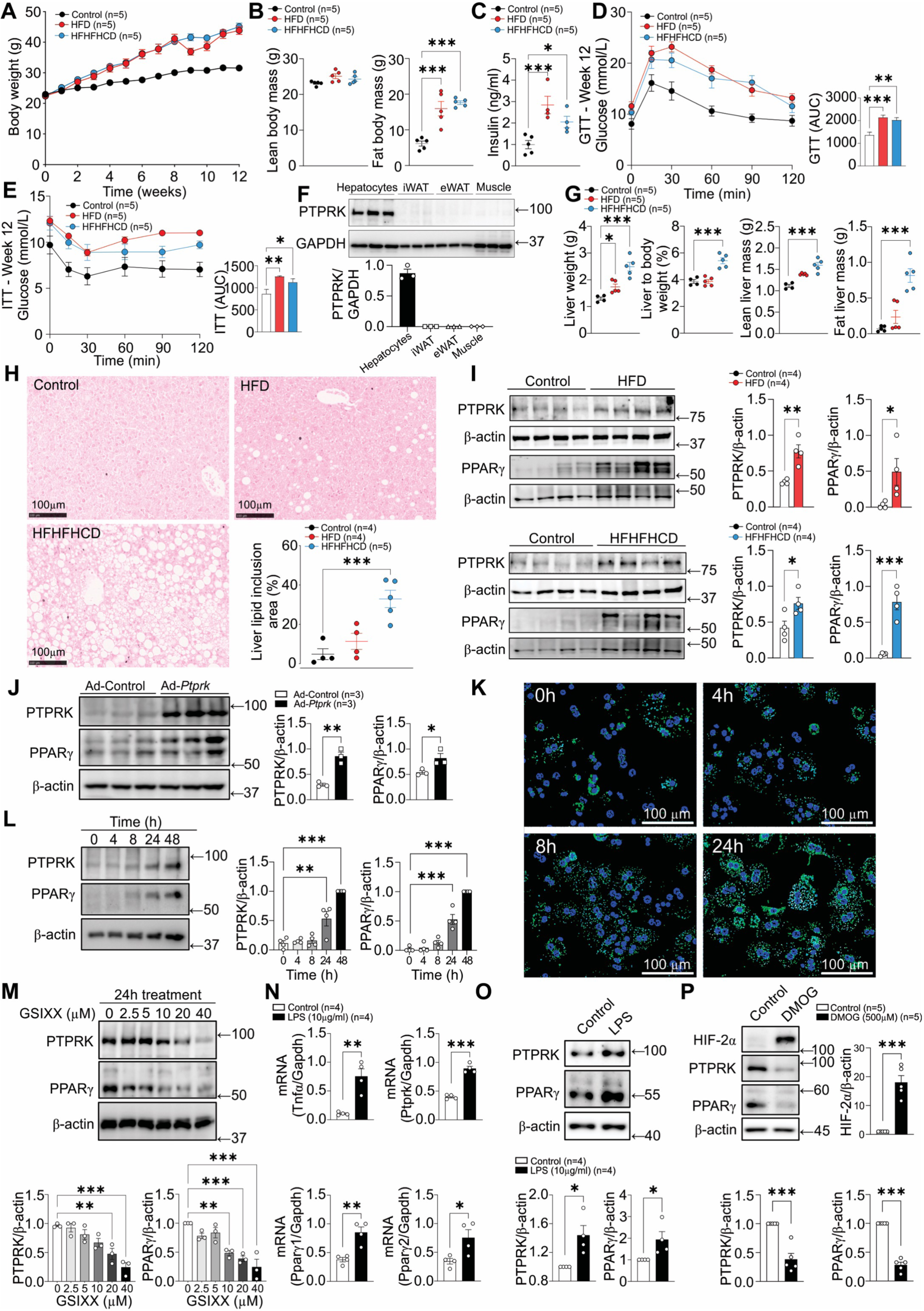
Hepatocyte PTPRK is induced by Notch signalling and LPS, correlating positively with PPARγ in obese mouse models and primary hepatocytes. (**A-E**). 8-week-old C57BL6N mice were fed either a high-fat diet (HFD) or a high-fat high-fructose high-cholesterol diet (HFHFHCD) for 12 weeks. We measured (**A**) body weight, (**B**) body composition, and (**C**) fasting insulinemia. Mice underwent (**D**) glucose and (**E**) insulin tolerance tests after 12 weeks of diet. (**F**) Primary hepatocytes, inguinal white adipose tissue (iWAT), epidydimal white adipose tissue (eWAT), and gastrocnemius (muscle) were harvested for immunoblot analysis of PTPRK. (**G**) The livers were extracted and assessed for liver weight and composition. (**H**) Histological analysis was conducted to quantify the vacuolation area, serving as an indicator of hepatic lipid inclusions. (**I**) Immunoblot analysis was carried out to determine the levels of PTPRK and PPARγ. (**J**) 8-week-old C57BL6N mice receiving a chow (control) diet were transduced with an adenoviral vector to induce PTPRK overexpression (Ad-*Ptprk*). Two weeks later, immunoblot analysis was performed in liver samples to assess the levels of PTPRK and PPARγ. (**K**) Primary mouse hepatocytes were cultured overnight under standard conditions and fixed at different time points (0, 4, 8, and 24h) for Nile Red staining to visualise lipid droplets. (**L**) Immunoblot analysis of PTPRK and PPARγ was performed on primary mouse hepatocytes collected at different time points as indicated. (**M**) Primary mouse hepatocytes were cultured overnight and treated with different concentrations of the Notch signalling inhibitor GSIXX for 24h. Immunoblot analysis was employed to evaluate the expression levels of PTPRK and PPARγ. (**N**, **O**) Primary mouse hepatocytes were cultured overnight and treated with lipopolysaccharide (LPS) for 24h. The gene expression was analysed by quantitative PCR (qPCR, **N**) or immunoblot techniques (**O**). (**P**) Primary mouse hepatocytes were cultured overnight and treated with dimethyloxalylglycine (DMOG), an inhibitor of 2-oxoglutarate-dependent dioxygenases required for hypoxia-inducible factor (HIF) degradation, for 24h. Immunoblot analysis was performed to examine the expression of HIF2α, PTPRK, and PPARγ. Statistical significance is denoted as \**p*<0.05, \*\**p*<0.01, \*\*\**p*<0.001.

Extended culture of primary mouse hepatocytes resulted in a loss of differentiation, causing changes in metabolic pathways similar to those observed *in vivo* during the progression of NAFLD/MASLD. We observed a gradual accumulation of lipid droplets in the cytosol of hepatocytes (**Fig. 2K**). This was accompanied with increased protein levels of PTPRK and PPARγ (**Fig. 2L**). Acute and chronic treatment of primary hepatocytes with insulin or pro-inflammatory cytokines TNFα, IL6 or IFNγ did not affect PTPRK expression (**Supplementary Fig. 2A-D**). NOTCH2 is significantly increased in primary hepatocytes over time in culture (**Supplementary Fig. 2E**), and the administration of the Notch signalling inhibitor GSIXX effectively prevented upregulation of both PTPRK and PPARγ in a dose-dependent manner (**Fig. 2M**). Treatment with lipopolysaccharide (LPS) significantly increased both PTPRK and PPARγ transcripts and protein levels (**Fig. 2N,O**). Additionally, primary mouse hepatocytes treated with dimethyloxalylglycine (DMOG), an inhibitor of 2- oxoglutarate-dependent dioxygenases, resulted in reduced PTPRK and PPARγ expression levels (**Fig. 2P**). Together, these experiments provided compelling evidence of a positive correlation between PTPRK and PPARγ in hepatocytes, which can be regulated by diverse signalling pathways.

### PTPRK deletion protects against diet-induced obesity, insulin resistance, and hepatic steatosis in mice

To directly evaluate the metabolic relevance of PTPRK, we conducted loss-of-function studies using 8-week-old *Ptprk*^-/-^ and *Ptprk*^+/+^ mice subjected to either an HFHFHCD or a chow diet. PTPRK deficiency had minimal impact on body weight gain and fat accumulation in chow-fed mice (**Supplementary Fig. 3A-D**). Glucose and insulin tolerance tests performed at 8 weeks of age showed no differences between *Ptprk*^-/-^ and *Ptprk*^+/+^ mice (**Supplementary Fig. 3E-H**). At 20 weeks, *Ptprk*^-/-^ mice showed improved glucose homeostasis (**Supplementary Fig. 3I-L**). The intake of chow diet was not affected (**Supplementary Fig. 3M,N**). After 12 weeks of HFHFHCD feeding, *Ptprk*^+/+^ mice developed obesity, characterised by substantial increases in body weight, fat mass, circulating insulin levels, homeostasis model assessment of insulin resistance (HOMA-IR), glucose intolerance and insulin resistance (**Fig. 3A-H, Supplementary Fig. 4A,B**). Strikingly, *Ptprk*^-/-^ mice displayed resistance to HFHFHCD-induced obesity, as their body weight, fat mass, circulating insulin levels, HOMA-IR, glucose sensitivity and insulin resistance were all significantly lower (**Fig. 3A-H**). This protective effect was particularly prominent in female mice.

**Figure 3.**
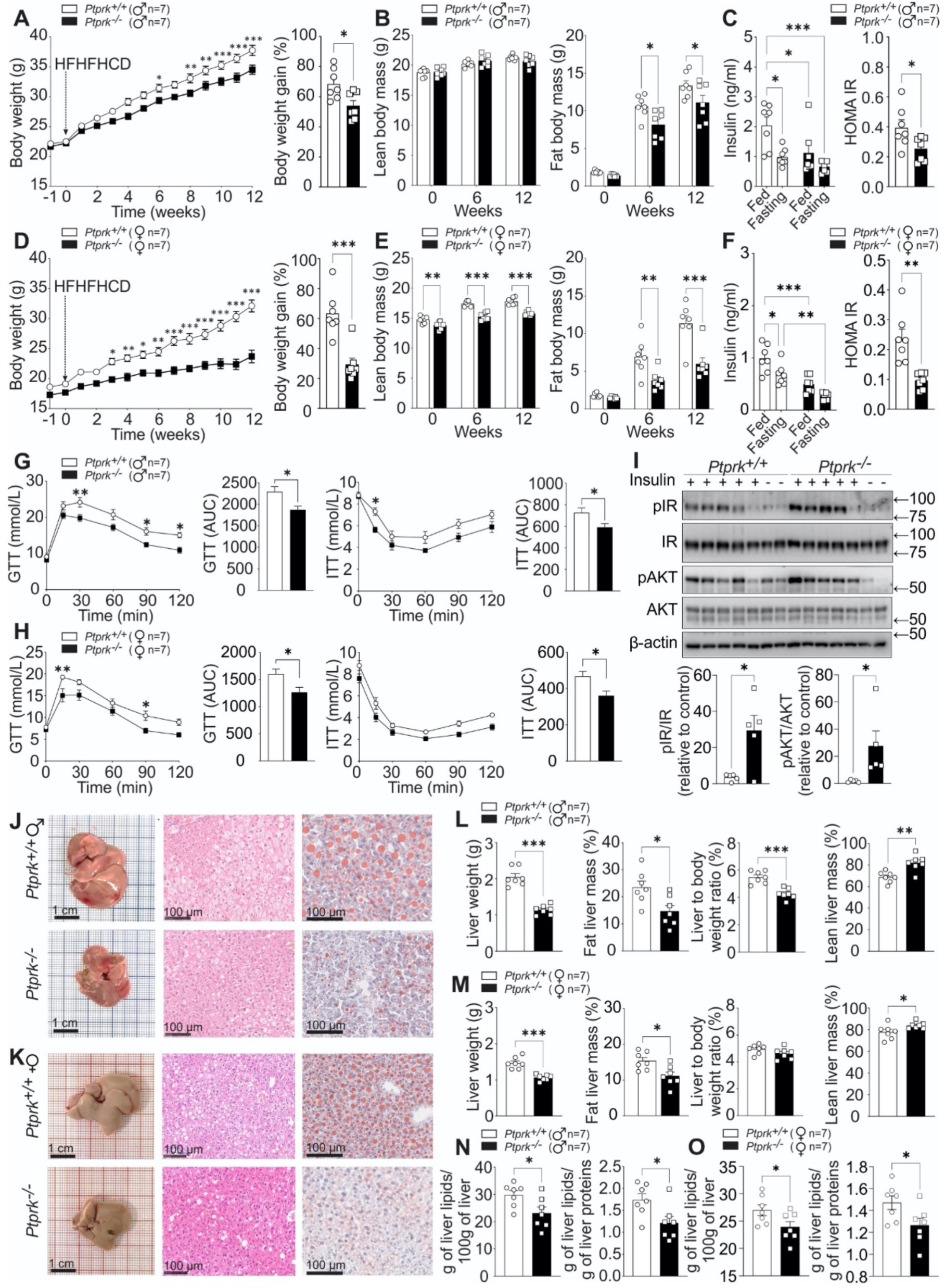
PTPRK deletion confers protection against diet-induced obesity, insulin resistance, and hepatic steatosis. (**A-H**) Male (♂) and female (♀) C57BL6N *Ptprk^+/+^*and *Ptprk^-/-^* mice, aged 8 weeks, were subjected to a high-fat, high-fructose, high-cholesterol diet (HFHFHCD) for a period of 12 weeks. We measured body weight (**A, D**), body composition (**B, E**), insulinemia (**C, F**), and performed glucose and insulin tolerance tests (**G, H**). (**I**) At the end of HFHFHCD feeding, insulin was administered to female mice 10 min prior to liver collection. Immunoblot analysis was employed to examine the expression of pIR and pAKT in the liver. (**J-O**) Liver samples were analysed to assess fat accumulation through histological examination (**J, K**), measurements of liver weight and composition (**L, M**), and total liver lipid extraction (**N, O**). (**I**). Statistical significance is denoted as \**p*<0.05, \*\**p*<0.01, \*\*\**p*<0.001.

**Figure 4.**
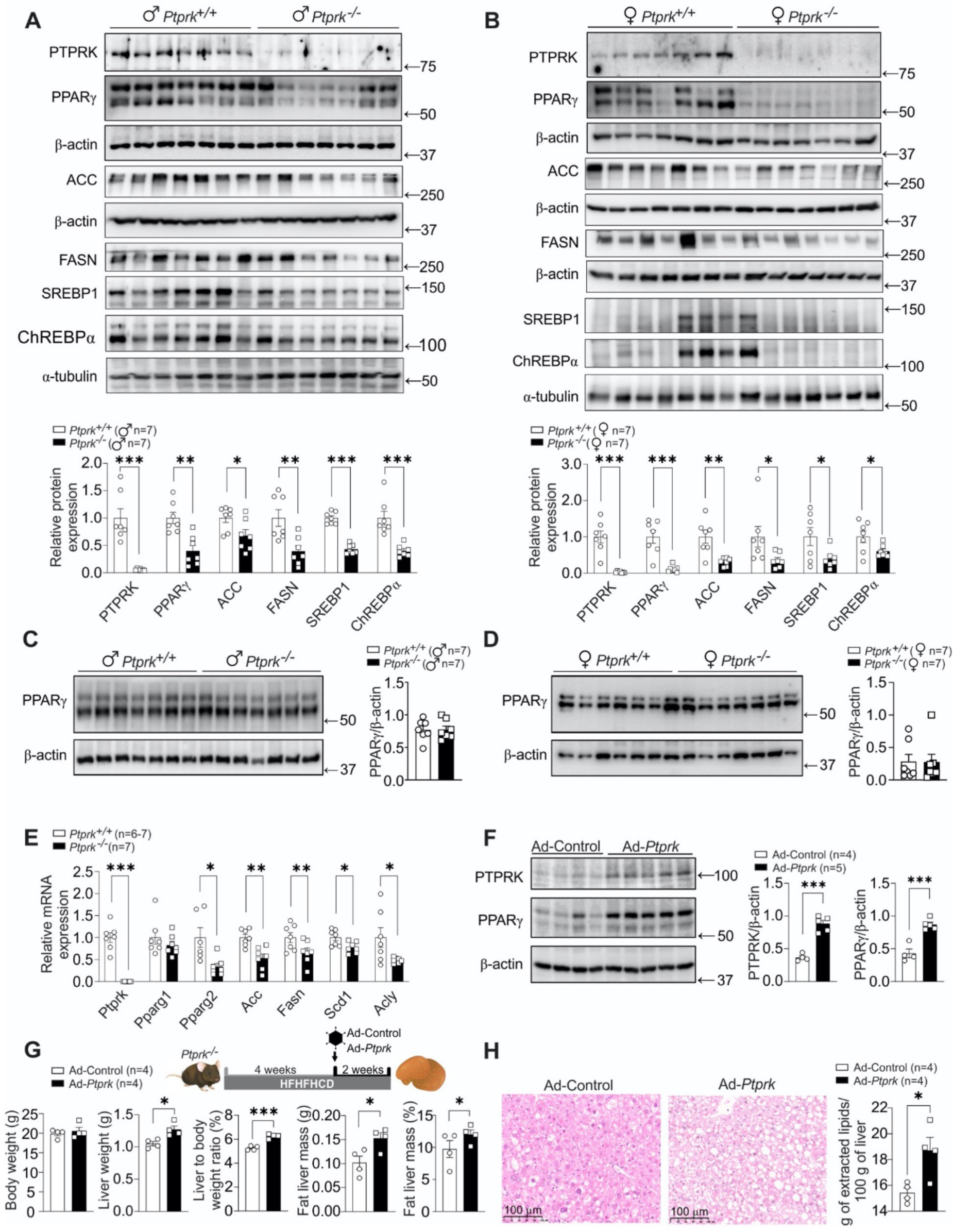
PTPRK orchestrates the hepatic expression of metabolic enzymes and transcription factors promoting steatosis in mice fed an obesogenic diet. (**A, B**) Eight-week-old male (♂) and female (♀) C57BL6N *Ptprk^+/+^* and *Ptprk^-/-^*were exposed to a high-fat, high-fructose, high-cholesterol diet (HFHFHCD) for 12 weeks. Liver samples were analysed to examine the levels of PTPRK, PPARγ, ACC (Acetyl-CoA Carboxylase), FASN (Fatty Acid Synthase), SREBP1 (Sterol Regulatory Element-Binding Protein 1), and ChREBP (Carbohydrate Response Element-Binding Protein). (**C, D**) Subcutaneous (inguinal fat, **C** and **D**) white adipose tissues were collected for immunoblot analysis of PPARγ. (**E**) Liver mRNA expression of Ptprk, Pparγ, Acc, Fasn, Scd1 (Stearoyl-CoA Desaturase 1), and Acly (ATP Citrate Lyase) was assessed. (**F**) Mice were subjected to HFHFHCD for four weeks and were administered an adenoviral vector to induce PTPRK overexpression (Ad-*Ptprk*). After a 2- week period, liver samples were collected for immunoblot analysis of PTPRK and PPARγ. (**G**) PTPRK knockout mice were subjected to HFHFHCD for four weeks and subsequently injected with Ad-*Ptprk*. After an additional two weeks on HFHFHCD, body weight was measured, and liver were collected for the evaluation of weight and composition. (**H**) Liver histological assessment and total lipid extraction were performed after PTPRK overexpression. Statistical significance is indicated as \**p*<0.05, \*\**p*<0.01, \*\*\**p*<0.001.

Consistent with the metabolic analyses, PTPRK-deficient mice exhibited elevated energy expenditure, specifically during the dark cycle (**Supplementary Fig. 4C**). *Ptprk^-/-^* mice also displayed increased VO2 levels during the night, and the respiratory exchange ratio (RER) showed a downward trend (**Supplementary Fig. 4C**). No significant disparities were noted in ambulatory activity or food and water intake (**Supplementary Fig. 4C,D**). The analysis of food intake over a span of 12 weeks revealed no disparities in males, but lower levels in female *Ptprk*^-/-^ mice, resulting in lower cumulative energy intake (**Supplementary Fig. 4E,F**). PTPRK deficiency did not result in altered lipid excretion through faeces (**Supplementary Fig. 4G**), suggesting that the reduced weight observed in *Ptprk*^-/-^ mice is not related to changes in intestinal fat absorption. *Ptprk*^-/-^ livers exhibited significantly higher induction of p-IR and p-AKT compared to *Ptprk*^+/+^ mice in response to insulin (**Fig. 3I**). The phosphorylation levels induced by insulin on IR and AKT displayed no discernible differences in primary hepatocytes (**Supplementary Fig. 5A**), indicating that PTPRK does not directly affect IR phosphorylation. We observed a significant reduction in hepatic lipid accumulation within the livers of *Ptprk*^-/-^ mice (**Fig. 3J-O**). Collectively, these findings demonstrate that while PTPRK-deficiency exerts minimal influence on normal development, it confers robust protection against diet-induced obesity, steatosis, and insulin resistance.

### PTPRK expression shapes nutrient-driven metabolic reprogramming in hepatocytes

Having established that PTPRK plays a major metabolic role in obesity, we next sought to define the lipogenic pathways affected in HFHFHCD-fed mice. Immunoblot analysis showed that *Ptprk*^-/-^ mice exhibited lower levels of hepatic PPARγ (**Fig. 4A, B**), while no differences were observed in subcutaneous and visceral adipose tissues (**Fig. 4C,D and Supplementary Fig. 5B,C**). We observed significantly reduced expression of Pparγ2 transcripts in *Ptprk*^-/-^ mice, while Pparγ1 was unaffected (**Fig. 4E**). Key lipogenic enzymes, namely Scd1, Acly, Acc, and Fasn mRNA were downregulated in *Ptprk*^-/-^ mice (**Fig. 4E**). Immunoblot analysis confirmed diminished levels of ACC and FASN in *Ptprk*^-/-^ mice (**Fig. 4A,B**). In addition, transcription factors governing fat metabolism, SREBP1c and ChREBP, also exhibited decreased expression in *Ptprk*^-/-^ livers (**Fig. 4A,B**).

Next, we used adenoviral-mediated upregulation of PTPRK in four week high-fat fed mice and observed a significant increase in hepatic PPARγ expression following adenoviral infection (**Fig. 4F**). Adenoviral PTPRK overexpression reverted the hepatic phenotype of obese *Ptprk^-/-^*mice, including increased liver weight, liver-to-body weight ratio, and liver fat mass (**Fig. 4G**). Histological examination of liver sections and lipid measurements revealed pronounced lipid deposition following PTPRK overexpression (**Fig. 4H**). These results demonstrate that hepatic PTPRK overexpression effectively reverses key phenotypic characteristics observed in PTPRK-deficient mice. Primary hepatocytes with reduced PTPRK levels (heterozygous) or complete deletion (knockouts) showed reduced kinetics of STAT1 phosphorylation in response to IFNγ (**Supplementary Fig. 5D,E**). STAT1 and Activator Protein 1 (AP-1) play a pivotal role in driving PPARγ expression and lipid accumulation within the liver^19,20^. Notably, we observed significantly lower levels of c-Fos/AP-1 in *Ptprk*^-/-^ livers (**Supplementary Fig. 5F**). Taken together, our results suggest that PTPRK acts upstream of transcriptional regulators of *de novo* lipogenesis and lipid metabolism in obesity.

### Phosphoproteomic analysis revealed FBP1 as a PTPRK substrate in hepatocytes during steatosis

To explore the mechanisms by which PTPRK inactivation in hepatocytes might drive the development of steatosis, we performed unbiased transcriptome and proteomic analysis. Hepatocytes were isolated and separated based on their fat content (**Fig. 5A**). Immunoblot analysis of high-fat content hepatocytes further established the positive correlation between PTPRK and PPARγ (**Fig. 5B**). Steatotic PTPRK-deficient hepatocytes have reduced Cd36 expression, a crucial PPARγ target in cellular fatty acid uptake (**Fig. 5C**). In contrast, Cpt1, facilitating long-chain fatty acid transport for mitochondrial β-oxidation, displayed an opposing pattern, with higher expression in *Ptprk*^-/-^ hepatocytes (**Fig. 5C**). We performed RNA-Seq analysis in low/high fat *Ptprk^-/-^*and *Ptprk^+/+^* hepatocytes (**Supplementary Fig. 6A**). Volcano plot analysis revealed that the predominant significant differences occurred among genes upregulated in low-fat hepatocytes compared to high-fat hepatocytes within the same genotype (**Supplementary Fig. 6B,C**). In contrast, only a limited number of genes exhibited significant transcriptional alterations resulting from PTPRK deletion in low-fat or high-fat hepatocytes (**Supplementary Fig. 6D,E**). We also observed reduced PPAR signalling in *Ptprk*^-/-^ hepatocytes (**Fig. 5D**). Comparison of low-fat to high-fat *Ptprk*^+/+^ hepatocytes revealed enriched pathways, including cell adhesion molecules, MAPK signalling, PI3K-AKT signalling, cytokine interaction, and chemokine signalling (**Fig. 5E**). In *Ptprk*^-/-^ hepatocytes, the same comparison highlighted pathways including gap junction, ECM receptor interaction, focal adhesion, cAMP signalling, PI3K-AKT signalling, and RAP1 signalling.

**Figure 5.**
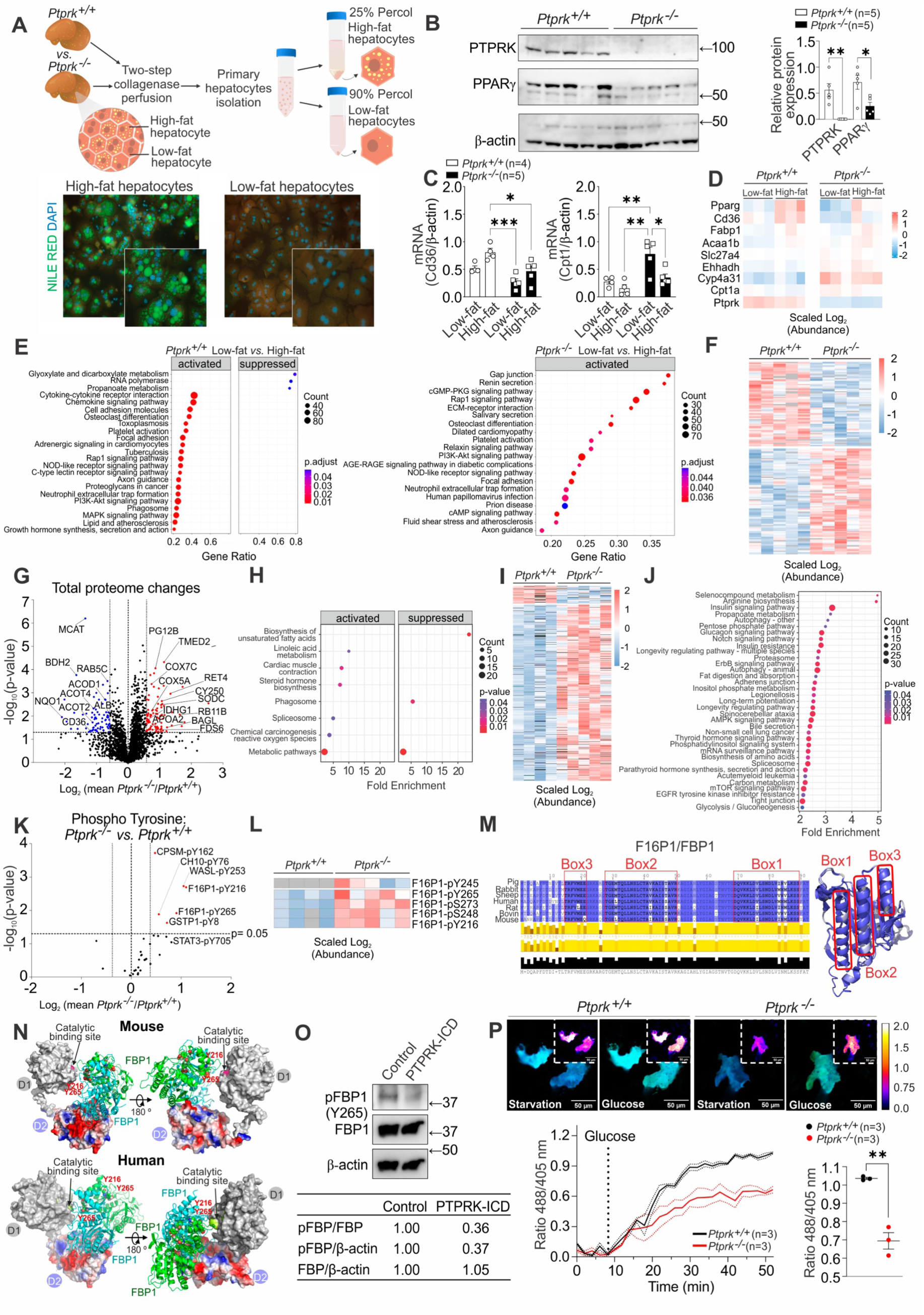
Comprehensive analysis of the transcriptome, proteome, and protein phosphorylation changes in primary hepatocytes isolated from livers of *Ptprk^-/-^* and *Ptprk^+/+^*mice. (**A**) Methodological approach schematic illustrating the isolation of primary hepatocytes from mice fed HFHFHCD for 12 weeks, followed by separation based on cell density into hepatocytes with high-fat content and hepatocytes with low-fat content. (**B**) Immunoblot analysis revealing PTPRK and PPARγ expression profiles of hepatocytes with high-fat content. (**C**) RT-qPCR analysis depicting changes in the expression of lipid metabolism-related genes. (**D**) RNA-Seq heatmap displaying alterations in PPAR pathway-related genes. (**E**) RNA-Seq KEGG pathway enrichment analysis comparing *Ptprk^+/+^* low-fat vs. high-fat hepatocytes (left side) and the same comparison in *Ptprk^-/-^* hepatocytes (right side). (**F**) Total proteome global heatmap showing significantly altered proteins. (**G**) Volcano plot illustrating the changes in the total proteomic profile between *Ptprk^-/-^*and *Ptprk^+/+^* high-fat hepatocytes. (**H**) Total proteome KEGG pathway enrichment analysis. (**I**) Phosphoproteome global heatmap showing significantly altered phosphoproteins. (**J**) Phosphoproteome KEGG pathway enrichment analysis. (**K**) Volcano plot displaying only the quantification of tyrosine phosphosites in *Ptprk^-/-^* and *Ptprk^+/+^* hepatocytes. Phosphosites with over 30% increase in *Ptprk^-/-^* cells are marked in red (p<0.05). (**L**) Heatmap showcasing the significantly changing phosphopeptides in fructose-1,6-bisphosphatase 1 (F16P1/FBP1). (**M**) Schematic representation of different F16P1/FBP1 amino acid sequences, indicating distinct boxes for interaction mapping experiments. The predicted helical regions are depicted in the three-dimensional structure on the right side. (**N**) Conservation mapping of the predicted PTPRK- FBP1 interface, illustrating the PTPRK-D2 complex (red, blue, and grey surface representation of their electrostatic surface potential) interacting with the FBP1 dimer (light green and light blue) and the proximity of the PTPRK catalytic site and increased FBP1 phosphotyrosine residues. The D1 domain of PTPRK is shown in grey surface representation. (**O**) Immunoblot analysis of pervanadate-treated mouse hepatocyte lysates incubated with or without the recombinant PTPRK-ICD (ICD: intracellular domain) prior to pFBP1 (Y265) analysis (**P**) Primary mouse hepatocytes were transfected with HYlight to monitor fructose 1,6-bisphosphate dynamics. After injection of 20 mM glucose, fluorescence ratios (R488/R405) were calculated and Min-Max normalized. Solid lines represent the mean across cells, while dots represent the mean±SEM. The presented data represent the average of multiple independent biological replicates. Statistical significance in panels **B**, **C**, and **P** is indicated as \**p*<0.05, \*\**p*<0.01, \*\*\**p*<0.001.

We next performed proteomics and phosphoproteomic analysis of hepatocytes with high-fat content (**Supplementary Fig. 6F**). The Venn diagram illustrates that 1148 proteins show modifications in both the phosphoproteomic and total proteome datasets. This suggests an intricate relationship between these protein datasets, indicating regulatory mechanisms acting at the translational level and post-translationally through phosphorylation (**Supplementary Fig. 6F**). The heatmap displays diverse protein changes between *Ptprk*^-/-^ and *Ptprk*^+/+^ hepatocytes (**Fig. 5F**), revealing their dynamic response. In *Ptprk*^-/-^ hepatocytes, an upregulation of specific proteins has been observed, reflecting a complex interplay of molecular events associated with altered mitochondrial function and redox balance, closely linked to cellular metabolic reprogramming (**Fig. 5G**). Enriched pathways include metabolism, phagosome, and biosynthesis of unsaturated fatty acids (**Fig. 5H**). PTPRK- deficiency increases phosphorylated residues across various proteins (**Fig. 5I**). These changes are associated with crucial pathways, including insulin signalling, mTOR pathway, AMPK signalling, insulin resistance, glucagon signalling, adherens junctions, and biosynthesis of amino acids (**Fig. 5J**).

A total of 2572 phosphosites were significantly upregulated in *Ptprk*^-/-^ hepatocytes compared with 258 found at lower levels (**Supplementary Fig. 6G**). Phosphotyrosine residues of CPSM (pY162), CH10 (pY76), WASL (pY253), GSTP1 (pY8), and FBP1 (pY265, pY216) were increased in *Ptprk*^-/-^ hepatocyte (**Fig. 5K**). The focussed analysis of FBP1, a hepatic tumour suppressor^21^, revealed changes also at the positions pS273, pS248, pY265, pY245 and pY216 in *Ptprk*^-/-^ steatotic hepatocytes (**Fig. 5L).** FBP1 is a key enzyme active in gluconeogenesis and glucose homeostasis. Structural modelling highlights conserved helical regions (**Fig. 5M**) that engage with the PTPRK D2 domain (**Fig. 5N**), placing phosphorylated tyrosine residues near the PTPRK catalytic D1 domain. Computational simulations confirmed PTPRK and tyrosine phosphorylated complex predictions with a range of different assemblies (**Supplementary Fig. 7A-F**). Pervanadate-treated hepatocyte lysates, combined with recombinant PTPRK intracellular domain (PTPRK-ICD), demonstrated FBP1 dephosphorylation (**Fig. 5O**). Liver analyses in female *Ptprk*^-/-^ mice following a 12-week HFHFHCD showed higher pFBP1 (pY265) levels (**Supplementary Fig. 8A**). We analysed glycolysis dynamics using HYlight, a biosensor designed to track real-time changes in intracellular levels of the FBP1 substrate, fructose 1,6-bisphosphate^22^. We observed a reduction in fructose 1,6-bisphosphate levels in *Ptprk*^-/-^ hepatocytes when stimulated with glucose (**Fig. 5P**). Our results demonstrate the dynamic interplay between PTPRK and FBP1, significantly impacting glucose metabolism.

### Deletion of PTPRK induces metabolic reprogramming in the liver during diet-induced obesity

To assess the importance of hepatic PTPRK in glycolytic control, we cultured primary mouse hepatocytes after adenovirus-mediated PTPRK overexpression or silencing (**Fig. 6A**). Glucose-starved hepatocytes overexpressing PTPRK displayed increased glycolytic activity. This was evident in the extracellular acidification rates measured after acute glucose injection and after oligomycin blockade of mitochondrial respiration (**Fig. 6B**). Elevated glycolysis channels pyruvate towards acetyl-CoA synthesis, triggering *de novo* lipogenesis. Hepatocytes with PTPRK overexpression exhibited increased lipid droplet accumulation (**Fig. 6C**). In addition, lipid droplet accumulation occurred to a greater extent in PTPRK overexpressing hepatocytes after fatty acid administration (**Fig. 6C**). Inhibition of glucose oxidation resulted in decreased PPARγ expression, while PTPRK was not affected (**Supplementary Fig. 8B**).

**Figure 6.**
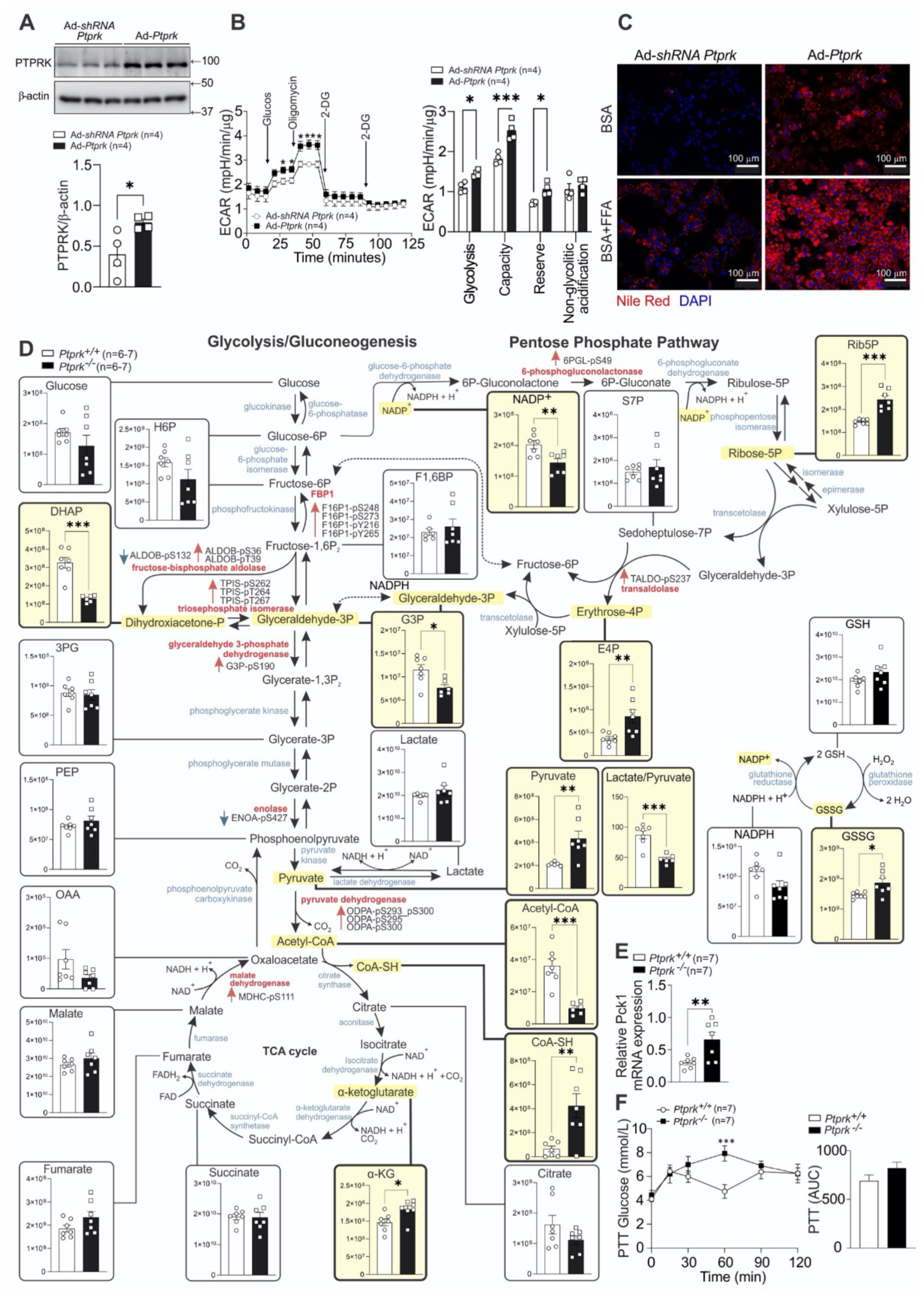
Hepatic PTPRK induces metabolic reprogramming in the livers of mice fed an obesogenic diet. (**A**) Primary mouse hepatocytes were cultured and transduced with an adenoviral vector to induce either PTPRK overexpression (Ad-*Ptprk*) or silencing of PTPRK (Ad-shRNA *Ptprk*). Immunoblotting (triplicates representative of 4 independent experiments) confirmed the modulation of PTPRK expression and (**B**) real-time measurement of extracellular acidification rate (ECAR) in response to glycolytic modulators, revealed changes in glycolytic parameters. (**C**) Primary mouse hepatocytes were treated with a mixture of BSA- conjugated fatty acids, palmitate (PA), and oleate (OA) (0.4 mM PA and 0.8 mM OA) to simulate triglyceride deposition, and subsequently stained with Nile Red to visualize lipid droplets rich in neutral lipids (triglycerides). (**D**) Untargeted metabolomics analysis was conducted in mouse livers from *Ptprk^-/-^*and *Ptprk^+/+^* female mice fed HFHFHCD for 12 weeks. Metabolites exhibiting statistically significant changes are highlighted (light yellow). The data is presented as raw abundances corrected for sample weight. Enzymes associated with these metabolites, which showed significant differences in the levels of phosphorylated amino acids based on the analysis presented in Figure 5I, are indicated in red. Red arrows near highlighted enzymes indicate increased phosphorylation at specific amino acid residues, while blue arrows indicate reduced phosphorylation at those respective residues. (**E**) qPCR analysis was performed to assess Pck1 mRNA levels in the livers of *Ptprk^-/-^* and *Ptprk^+/+^* mice. (**F**) A pyruvate tolerance test was conducted in mice fed HFHFHCD for 12 weeks after overnight fasting to assess their gluconeogenic capacity in response to pyruvate administration. Statistical significance is indicated as \**p*<0.05, \*\**p*<0.01, \*\*\**p*<0.001.

We next sought to validate our results in human hepatocytes. *PTPRK^-/-^* and *PTPRK^+/+^* human embryonic stem cells (hESC) were differentiated into hepatocyte-like cells (HLCs, **Supplementary Fig. 8C-E**). Deletion of PTPRK did not affect hESC hepatocyte differentiation nor the ability of HLCs to produce and secrete albumin (**Supplementary Fig. 8C-E**). Similar to mouse hepatocytes, PTPRK-deficient HLCs exhibited a reduced glycolytic rate following glucose stimulation (**Supplementary Fig. 8F**). Together, these observations indicate that PTPRK leads to steatosis indirectly by stimulating glycolytic activity and directly by accelerating fatty acid esterification and lipid droplet formation in response to fatty acids.

Liver metabolites from *Ptprk*^-/-^ and *Ptprk*^+/+^ mice fed HFHFHCD for 12 weeks were quantified by mass spectrometry (**Fig. 6D**). *Ptprk*^-/-^ livers showed decreased levels of dihydroxyacetone phosphate and glyceraldehyde-3-phosphate, with a corresponding reduction in the lactate/pyruvate ratio. *Ptprk*^-/-^ livers displayed elevated α-ketoglutarate levels and increased pyruvate. Despite elevated pyruvate levels, *Ptprk*^-/-^ livers exhibited reduced concentrations of acetyl-CoA, and increased free coenzyme A compared to *Ptprk*^+/+^. This aligns with our findings of decreased levels of pyruvate dehydrogenase phosphatase in *Ptprk*^-/-^ mice, while no differences were observed for pyruvate dehydrogenase kinase (**Supplementary Fig. 8G**). PTPRK deficiency also led to heightened pentose phosphate pathway (PPP) intermediates, particularly ribulose-5-phosphate and erythrose 4-phosphate (**Fig. 6D**). Enhancing the flux through the PPP could reinforce essential production of reducing equivalents, and increase oxidative stress management. Parallel to shifts in lactate-to-pyruvate ratio, *Ptprk*^-/-^ livers unveiled elevated GSSG (**Fig. 6D**) and methionine sulfoxide levels (**Supplementary Fig. 8H**), indicative of an oxidised environment. The ratio of the classical redox indicators NAD^+^/NADH, NADP^+^/NADPH, and GSSG/GSH showed no significant changes (**Supplementary Fig. 8I**), although the total levels of NADP^+^ were significantly lower in *Ptprk*^-/-^(**Fig. 6D**). No differences were found in phosphorylated adenine nucleotides (ATP, ADP and AMP, **Supplementary Fig. 8J**) and amino acids (**Supplementary Fig. 8K**). We observed heightened expression of Pck1, a pivotal gluconeogenic driver, in *Ptprk*^-/-^ livers (**Fig. 6E**), consistent with lower glycolysis. *Ptprk*^-/-^ female mice, subjected to a 12-week HFHFHCD obesogenic diet, exhibited elevated blood glucose levels compared to *Ptprk*^+/+^ after pyruvate injection, supporting a shift to a more gluconeogenic state upon PTPRK deletion (**Fig. 6F**). Taken together, PTPRK plays a crucial role in controlling liver metabolism through the regulation of glycolytic intermediates and altered lipid dynamics.

### PTPRK contributes to hepatocyte transformation in obesity-associated HCC

Glycolytic and gluconeogenic proteins, including FBP1 (**Supplementary Fig. 9A**), contribute to HCC development^21^. We observed a stratification pattern based on PTPRK mRNA expression levels in human samples (**Fig. 7A**) that bifurcated into two distinct clusters: one characterised by high PTPRK expression and the other marked by low PTPRK expression (**Fig. 7B**). Normal liver samples uniformly exhibited low PTPRK expression, while in the context of NASH, peritumour, and tumour conditions, high PTPRK expression was correlated with elevated hepatic expression of glycolytic genes. The analysis of all liver samples and the focused analysis of tumour samples revealed a positive correlation between elevated PTPRK expression and hepatic expression of lipogenic genes (**Fig. 7B**). The enriched pathways associated with elevated PTPRK expression in liver tumour samples, as defined through KEGG pathway enrichment analysis, underscored the activation of fatty acid metabolism, type 1 diabetes mellitus, glycolysis/gluconeogenesis, TCA cycle, primary bile acid biosynthesis, biosynthesis of unsaturated fatty acids, PPAR signalling pathway, steroid biosynthesis, and oxidative phosphorylation (**Fig. 7C**).

**Figure 7.**
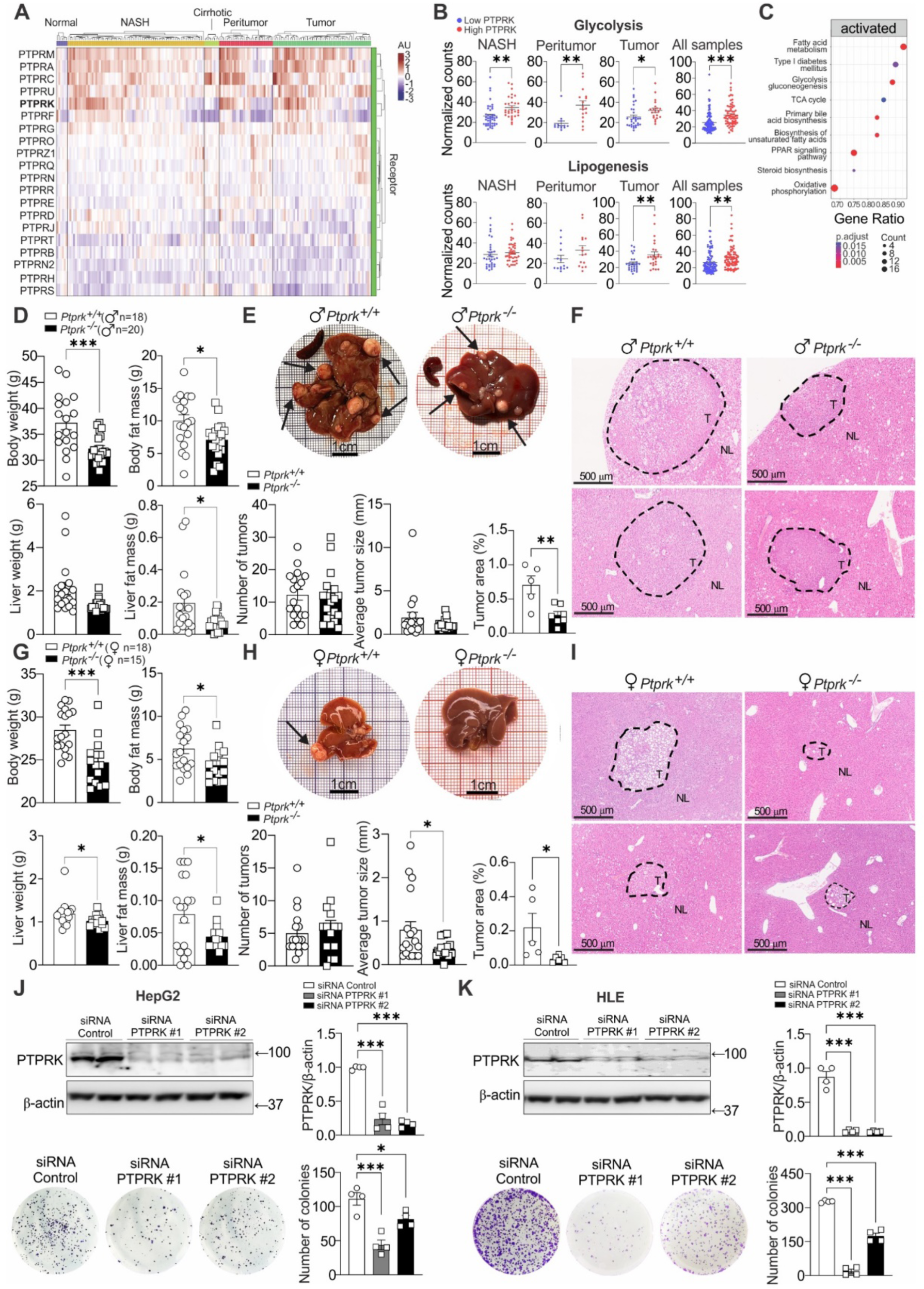
Influence of PTPRK in hepatocellular carcinoma (HCC) development. (**A**) RPTP mRNA expression profiling was conducted on human livers from dataset GSE164760 encompassing various stages of obesity-associated liver dysfunction and hepatocarcinogenesis, including normal liver, non-alcoholic steatohepatitis (NASH), cirrhotic livers, peritumour regions, and hepatocellular carcinoma (HCC) tumours. (**B**) Based on the expression levels of PTPRK, the samples were categorized as high or low, and the normalized counts of genes involved in glycolysis/gluconeogenesis and lipogenesis were analysed. (**C**) KEGG pathway enrichment analysis was performed specifically on tumour samples with low or high PTPRK expression levels. (**D-I**) Male and female *Ptprk^-/-^* and *Ptprk^+/+^*mice were subjected to diethylnitrosamine (DEN) induction of liver cancer at 2 weeks of age. Tumour development was assessed when the animals reached 40 weeks of age. Measurements of body weight, fat body mass, liver weight, and fat liver mass were recorded (**D, G**). Tumours on the hepatic lobes were quantified and measured, considering tumours bigger than 0.2mm. The results are presented as the number of tumours per liver and the average tumour size (**E, H**). Microscopic tumours were quantified through histological analysis. Representative H&E- stained sections showing nodules (**F, I**). (**J, K**) Human hepatoma cell lines HepG2 (**J**) and HLE (**K**) were transfected with siRNAs targeting PTPRK or siRNA control, and colony-forming capacity was assessed. Immunoblot analysis confirmed the efficiency of transfection, and crystal violet staining was employed to visualise and quantify the colonies. Statistical significance is indicated as \**p*<0.05, \*\**p*<0.01, \*\*\**p*<0.001.

To investigate the implications of PTPRK deletion in the context of liver cancer, diethylnitrosamine (DEN), a potent hepatocarcinogen, was administered by a single injection into *Ptprk*^-/-^ and *Ptprk*^+/+^ mice at the age of two weeks. *Ptprk*^-/-^ male mice showed reduced body fat accumulation, while *Ptprk*^-/-^ females exhibited diminished body weight and fat accumulation at the end of the experimental timeline compared with *Ptprk*^+/+^ (**Fig. 7D,G**). Livers from *Ptprk*^-/-^ male and female mice were smaller, with reduced absolute hepatic lipid content, although the relative liver fat content was unaltered (**Fig. 7D,G**). PTPRK deficiency did not affect DEN-mediated tumour formation (**Fig. 7E,H**). However, macroscopic evaluation showed reduced tumour size in *Ptprk*^-/-^ females (**Fig. 7H**). This was confirmed by histological analysis of liver sections, where *Ptprk*^-/-^ tumours exhibited significantly diminished dimensions and reduced fat accumulation regardless of gender (**Fig. 7E,F and 7H,I**). In line with these findings, silencing of PTPRK using siRNA in HepG2, HLE, and Huh6 cell lines (**Fig. 7J,K and Supplementary Fig. 9B**) led to a substantial attenuation of colony-forming capacity. Overall, our experiments support an oncogenic role of PTPRK in promoting rapid hepatic tumour growth.

## Discussion

Modern diets and lifestyles pose a chronic challenge to the ancestral mechanisms designed to control energy balance in humans. In the present study, we observed increased expression of hepatic RPTPs in steatosis and NASH, suggesting an adaptive response. The upregulation of PTPRK may serve as a part of a compensatory mechanism to mitigate the impact of disrupted cell adhesion components and mechanotransduction in fat-loaded hepatocytes. PTPRK seems to play a pivotal role in disease progression, influencing glycolysis, *de novo* lipogenesis signalling, and associated metabolic pathways.

We demonstrated that the upregulation of hepatocyte PTPRK-PPARγ correlates with NOTCH2 activation and that the inhibition of Notch signalling suppresses PTPRK-PPARγ expression in hepatocytes^17,23^. In addition, LPS boosts PTPRK-PPARγ, while HIF2α represses it. These results suggest that inflammation and hypoxia influence PTPRK expression, potentially contributing to the differences observed in human liver samples. The correlation between elevated PTPRK expression and the activation of glycolytic and lipogenic genes is consistent with the role played by PTPRK in regulating these metabolic pathways.

Obesogenic diets lead to hepatic PTPRK overexpression, and global PTPRK knockout mice are resistant to diet-induced metabolic dysfunction. The absence of PTPRK resulted in delayed onset of obesity and NAFLD/MASLD with suppression of nutrient-sensitive transcription factors PPARγ, SREBP1c, and ChREBP, relevant for metabolic reprogramming^24,25^. We found that PTPRK deletion leads to c-Fos downregulation and lower STAT1 activation in response to IFNγ. Both c-Fos and STAT1 are known to promote PPARγ expression^19,20^. Interestingly, we demonstrated that inhibition of glucose oxidation suppresses PPARγ expression in hepatocytes without altering PTPRK expression. Thus, PTPRK-induced glycolysis can contribute to increased PPARγ expression. Glucose oxidation provides the energy, metabolites, and reducing agents necessary for the execution of *de novo* lipogenesis and sustains a shift in fatty acid metabolism from a catabolic to an anabolic direction^26^. PTPRK deficiency results in lower expression of the PPARγ target gene CD36, a long-chain free fatty acid transporter that promotes the growth of HCC cells by increasing glycolysis^27^. In overnutrition, PTPRK mediates the shift towards glycolysis and fat storage. PTPRK deletion also protected mice against insulin resistance and hyperinsulinemia after obesogenic diet feeding. Hepatic PTPRK overexpression is sufficient to reverse the phenotype. Elevated insulin levels, insulin resistance, and excessive fat accumulation in hepatocytes are key drivers of metabolic reprogramming, fuelling the phenotypic changes necessary for malignant transformation of hepatocytes^4,28^.

In aerobic conditions, pyruvate and cytosolic NADH are readily oxidised in the mitochondria. However, as mitochondrial energy production meets cellular energy demand, counterregulatory mechanisms are activated to slow down the TCA cycle and mitochondrial function^29^. If the glycolytic activity remains active, an excessive supply of acetyl-CoA originating from pyruvate may accumulate, along with increased pyruvate fermentation into lactate^30^. The accumulated acetyl-CoA is then converted to malonyl-CoA, serving as a substrate for *de novo* lipogenesis and an allosteric inhibitor of CPT1α^26^. We found that the metabolic balance is regulated by PTPRK and favoured the conversion of pyruvate into acetyl-CoA by the pyruvate dehydrogenase complex. Indeed, we observed PTPRK-dependent increased lactate/pyruvate ratio, reduced PDHK1/PDP1 ratio, accumulation of acetyl-CoA, and upregulation of ACC and FASN. All these factors collectively contribute to the increased lipogenic capacity of PTPRK-expressing hepatocytes, ultimately resulting in higher steatosis.

Glycolytic and gluconeogenic rates are reciprocally regulated, and the suppression of gluconeogenic reactions favours glycolysis. The involvement of PTPRK in hepatic glucose metabolism is an important determinant of the metabolic phenotype of PTPRK knockout mice and the outcome of liver tumour formation experiments. Discrepancies in the role of PTPRK in tumour development and growth in extrahepatic tissues^31^ may be due to the PTPRK–FBP1 regulation and its hepatic role in gluconeogenesis. Our experiments revealed that pyruvate and α-ketoglutarate are elevated in PTPRK knockout mice. α-ketoglutarate is a TCA cycle metabolite, that can supply carbons for gluconeogenesis; however, its accumulation reduces liver gluconeogenesis^32^ and is associated with cancer suppression mechanisms^33^. The build-up of these gluconeogenic substrates indicates that even with a higher gluconeogenic capacity, the livers of PTPRK knockout mice still possess regulatory mechanisms to prevent uncontrolled gluconeogenesis. In line with this interpretation, hyperglycaemia was not observed in PTPRK knockout mice, but higher glucose production was found during the pyruvate tolerance test, in which animals were fasted to stimulate gluconeogenesis.

Fat-loaded hepatocytes isolated from PTPRK knockout mice showed an accumulation of phosphorylated FBP1, and real-time detection of the metabolite fructose 1,6-bisphosphate in primary hepatocytes lacking PTPRK revealed lower levels after glucose stimulation. Lower fructose 1,6-bisphosphate levels during glycolysis may also be a direct implication of the FBP1 tyrosine phosphorylation status. Thus, the observed outcome implies a futile cycle between FBP1 and PFK1 activities during glycolysis and optimised glucose production during gluconeogenesis. The substrate cycling at this step of glycolysis leads to wasteful ATP consumption^34^, in agreement with the observed increased energy expenditure in PTPRK knockout mice and the substantial disparities observed for fat accumulation despite minor differences in food intake. Further research is needed to explore the precise implications of tyrosine phosphorylation on FBP1. Additionally, it is crucial to identify the tyrosine kinases responsible for phosphorylating FBP1 and to examine the potential role of substrate cycling in glycolysis in maintaining energy homeostasis.

Besides PTPRK, several classical PTPs present promising therapeutic opportunities. However, developing selective and bioavailable PTP inhibitors has proved challenging^35^. Recent studies have shown the effectiveness of competitive inhibitors for closely related PTPN2 and PTPN1^36,37^. RPTPs can additionally be inhibited by inducing their dimerization. For example, antibodies targeting PTPRD ectodomains induce protein dimerization and degradation, thereby suppressing PTPRD-dependent cell invasion in a metastatic breast cancer cell line^38^. Therapeutic options for end-stage liver diseases are limited^39^. PTPRK inhibitors may hold potential in high-expressing PTPRK livers, as an alternative or adjuvant treatment for obesity-associated liver dysfunction.

In conclusion, PTPRK expression is increased in diseased human and mouse livers. Elevated hepatic PTPRK expression triggers heightened glycolysis, culminating in the activation of PPARγ and the stimulation of *de novo* lipogenesis. In mice, genetic PTPRK inhibition offers protection against the rapid development of liver dysfunction associated with obesogenic diet. Therefore, PTPRK emerges as a dual role player – serving as a biomarker for hepatic metabolic adaptations that influence the risk of metabolic liver disease and as a target for the development of new therapies. Screening and stratification of patients with NAFLD/MASLD based on hepatic PTPRK expression levels could guide therapeutic decisions to attenuate metabolic dysfunction associated with obesity.

## Materials and Methods

### Reagents

Human insulin solution, sodium pyruvate, sodium palmitate, oleic acid, bovine serum albumin, 2-deoxy-D-glucose, D-glucose, mannoheptulose, DMOG, and LPS were obtained from Sigma-Aldrich. Recombinant murine IFNγ (315-05-100ug PeproTech), recombinant human IL6 (206-IL-010, R&D Systems), insulin ProZinc (NDC 0010-4499-01, Boehringer Ingelheim, Rhein, Germany) were used for *in vivo* experiments.

### Human samples

We studied 19 biopsy specimens of patients undergoing a liver biopsy for medical reasons. The clinical characteristics of these patients are shown in **Supplementary Table 1**. Biopsies were collected after approval of the Hôpital Erasme Ethics Committee. Written informed consent was obtained from each participant.

### Extraction of proteins from human liver biopsies, enrichment of PTPs, and proteomics analysis

Frozen human liver biopsies were subjected to disruption using beads beating and sonication. Lysates were treated with lysis buffer containing 10% glycerol, 1% NP-40, cOmplete™EDTA-free protease inhibitor cocktail (Roche Diagnostics), and 1× phosphatase inhibitor (Sigma-Aldrich). After sonication, centrifugation at 20,000g for 1h at 4°C separated insoluble debris, retaining the supernatant for total proteome analysis. The obtained lysates were enzymatically digested using trypsin, targeting C-terminal lysine and arginine residues, except when adjacent to a C-terminal proline. Purification of the resulting peptides was performed using reverse-phase Sep-Pak C-18 cartridges, removing salts and buffers. By employing a strategy that explores the oxidation of cysteine in the catalytic site of PTPs, peptides containing cysteine residues within the PTP signature motif HCX5R were enriched through immunoprecipitation. Immunoprecipitated peptides, resuspended in 0.2% formic acid, were injected in triplicate for LC-MS/MS analysis. A 40-min reverse-phase gradient separation on UHPLC 1290 (Agilent Technologies) was followed by analysis on an Orbitrap Q Exactive HF mass spectrometer (Thermo Fisher Scientific), with MS scans spanning the 375–1500m/z range at 60,000 resolution. Data were acquired in Data-Dependent Acquisition mode, selecting the top 7 precursor ions for HCD fragmentation, followed by MS/MS analysis at 30,000 resolution. MaxQuant (version 2.0.3.0) processed spectral files, searching the Homo sapiens Uniprot database with FDR restricted to 1%.

### Mice

Mice were housed and managed in compliance with the Belgian Regulations for Animal Care, and the animal protocols underwent approval from the Commision d’Ethicque du Bien-Être Animal (CEBEA), Faculté de Médecine, Université libre de Bruxelles (dossier No. 732). Animals were housed at 22°C on a 12:12-h light-dark cycle with ad libitum access to food and water. *Ptprk* knockout mice were generated at The Jackson Laboratory (Ptprk-8356J-M669 project) by CRISPR/Cas9 technology and were bred on a pure C57BL/6N background. The strategy involved an intragenic deletion spanning 555 base pairs on Chromosome 10. This genetic alteration led to the excision of exon 3 within the *Ptprk* gene, accompanied by the removal of 283 base pairs from adjacent intronic sequences. The resulting mutation is predicted to induce an alteration in the amino acid sequence following residue 74 and an early truncation by 2 amino acids.

By breeding *Ptprk*^+/-^ mice we obtained *Ptprk*^−/−^ and *Ptprk*^+/+^ males and females littermates. *Ptprk*^+/+^ and *Ptprk*^-/-^ mice, aged 8 weeks, were randomly assigned to experimental diet-induced obesity feeding with unrestricted access to the specific diets: a HFD (60 kcal% fat D09100310i), a HFHFHCD (40 kcal% Fat, 20 kcal% Fructose, and 2% Cholesterol, D09100310i), or a control diet (10 kcal% Fat, D09100304i) from Research Diets (New Brunswick, NJ, USA). The duration for which the animals were subjected to the experimental diets ranged from 4 to 24 weeks, as indicated.

### Metabolic analysis

Evaluation of body and liver lean and fat mass was performed with EchoMRI™ 3-in-1 (NMR) body composition analyser from EchoMedical Systems (Houston, TX, USA).

Glucose tolerance tests were performed in 6h fasted mice with an intraperitoneal administration of glucose (2g D-Glucose/kg body weight). For pyruvate tolerance tests, mice were fasted overnight and administered pyruvate (2g/kg). Insulin tolerance tests were performed on mice fasted for 4h, with an intraperitoneal injection of insulin (0.75U/kg body weight). Fresh D-glucose, pyruvate, or insulin solutions were prepared in PBS immediately before the injections. Blood samples were obtained from the tail tip, and glycemia was measured using a glucometer (Accu-Check Performa, Roche, Basel, Switzerland). Blood serum was collected in a fed state (9am) or 6h after fasting (3pm) and insulin levels measured by ELISA (Crystal Chem Inc.).

At 18 weeks of age, *Ptprk*^−/−^ and *Ptprk*^+/+^ mice fed HFHFHCD for 10 weeks, were placed in metabolic cages TSE Phenomaster setup (TSE, Germany) for a duration of 72h. Following a 24h period of acclimatization, metabolic parameters, including physical activity, energy expenditure, and substrate utilization were assessed by indirect calorimetry.

### DEN-induced HCC

Liver tumour formation was induced by administering 25mg/kg of DEN in PBS via intraperitoneal injection into the underbelly region of 14-day-old mice. The mice were maintained on a chow diet. At 40 weeks of age, the mice were euthanized through cervical dislocation, and their livers were extracted for comprehensive analysis, including macroscopic and histological assessment of tumour number and size.

### Histological analysis

Mouse liver tissues intended for histological analysis were collected from euthanized mice, dissected, and subsequently rinsed with PBS. The tissues were fixed in 4% buffered formaldehyde (pH 7.4), embedded in paraffin blocks, sectioned into slices measuring 5-7µm using a Leica rotator microtome and stained with haematoxylin and eosin (H&E).

For immunohistochemistry analysis of PTPRK in human liver samples, 7µm thick paraffin sections were situated on positively charged slides. Antigen unmasking was performed with a heated citrate buffer (10mM, pH6.0). The sections were permeabilized using triton (0.1%), subsequently blocked with 2% milk, and incubated with 10% normal goat serum to prevent nonspecific binding. Primary antibodies were incubated overnight at 4°C, followed by incubation with goat anti-rabbit horseradish peroxidase secondary antibody (P044801). Negative controls were established by subjecting specimen slices solely to the secondary antibody.

Lipid accumulation was assessed by Nile Red staining (Sigma-Aldrich N3013). Primary hepatocytes were isolated and seeded onto chambered coverslips (IBIDI, 80806) at a density of 50,000 cells per well, 4h before adenoviral transfection for either PTPRK overexpression or silencing. After 24h of transfection, the culture medium was replaced with BSA-conjugated free fatty acids (sodium palmitate 0.4mmol/L, oleic acid 0.8mmol/L) or free fatty acid-free 1% BSA control-enriched medium with 1% FBS. The hepatocytes were fixed in 4% formaldehyde solution for 20min and stained with a 5μg/mL Nile Red solution and the nucleus were stained with DAPI. The stained cells were observed using an inverted fluorescence microscope (Axio Observer D1, Carl Zeiss, Oberkochen, Germany). The same staining procedure was applied to hepatocytes with high-fat and low-fat content, fixed and stained following an overnight culture period.

Hepatic lipid content was assessed in frozen sections of both *Ptprk*^+/+^ and *Ptprk*^−/−^ livers through Oil Red O (ORO) (Sigma-Aldrich, O1391) staining. Liver sections from cryostat cuts were equilibrated for 30min at room temperature in laminar flow hood. ORO working solution (0.3% ORO in 60% isopropanol) was applied to ensure complete coverage and incubated at 37°C. After counterstaining with haematoxylin, images were captured using NanoZoomer Digital Pathology (Hamamatsu Photonics K.K., version SQ 1.0.9) at 40x magnification.

### Lipid extraction

Total hepatic lipid content was evaluated by gravimetry after lipid extraction. Livers were removed, immediately freeze-clamped in liquid nitrogen, and stored at −80°C. The liver samples (100mg) were homogenized using a bead tissue homogenizer with cold methanol. After sonication, the homogenate was transferred to Falcon tubes. Chloroform was added, and the mixture was vortexed. Following agitation overnight at 4°C, the samples were centrifuged at 13,500g for 10min. The organic phase was collected, allowed to air-dry at room temperature, and the resulting pellet was weighed for total fat quantification. The solid middle layer formed during centrifugation was dried and weighed to determine the protein content. The results were expressed as mg of fat/100g of liver and g of fat/g of protein in the liver.

### LC-MS analysis of glycolytic, tricarboxylic acid (TCA) cycle, and pentose phosphate pathway (PPP) intermediates

The metabolomics analysis was conducted at the VIB Metabolomics Core (Leuven, Belgium). Polar metabolites were extracted using a two-phase methanol-water-chloroform method^40^. Dried metabolite samples were reconstituted in a solution of 60% acetonitrile and then transferred to LC-MS vials. For the analysis, an UltiMate 3000 LC System (Thermo Scientific) was coupled to a Q-Exactive Orbitrap mass spectrometer. Separation was achieved using a SeQuant ZIC/ pHILIC Polymeric column (Merck Millipore). A gradient of solvent A (95% acetonitrile-H2O, 2 mM ammonium acetate pH 9.3) and solvent B (2 mM ammonium acetate pH 9.3) was employed. Mass spectrometry was performed in the negative ion mode, encompassing both full scans and a targeted Selected Ion Monitoring (SIM) approach. Data acquisition was managed using Xcalibur software (Thermo Fisher Scientific)^41^. The data is presented as raw abundances corrected for sample weight.

### Primary mouse hepatocyte isolation, cell culture and treatments

Mouse primary hepatocytes were isolated from *Ptprk*^-/-^ and *Ptprk*^+/+^ mice following overnight ad libitum feeding, utilizing a two-step collagenase perfusion method through the vena cava. The process was initiated by anaesthetizing the mice through an intraperitoneal injection of a ketamine (100mg/kg) and xylazine (10mg/kg) mixture, peritoneum was opened, and the infra-hepatic segment of the vena cava was cannulated for perfusion. The portal vein was cut to clear blood from the liver. In the first perfusion step, the liver was exposed to HBSS (Thermo Fisher Scientific) supplemented with 10mM HEPES (pH 7.4), saturated with O2/CO2 (95:5 vol/vol), at 37 °C for 10min. The second step involved adding collagenase type IV (0.3 mg/mL) to William’s E Medium (Thermo Fisher Scientific) and further perfusing for 10min, effectively digesting the liver tissue. The digested liver was transferred to a sterile plastic dish, and cells were dispersed using a coarse-toothed comb in cold William’s E Medium, followed by filtration through a 100-µm cell filter to eliminate cell clumps. The resulting clump-free cell suspension was pelleted through centrifugation at 50g for 5min at 4 °C, the pellet was resuspended in William’s E Medium and layered onto Percoll^®^ solution (Sigma-Aldrich) (10ml Percoll^®^+ 1.25ml PBS 10X + 1.25 ml H2O.) and centrifuged for 10 min at 1000 RPM. The pellet was resuspended in William’s E Medium and washed 3 times (centrifugation at 50g for 5min at 4 °C). Viability assessment using the trypan blue exclusion test yielded approximately 15-20 million cells with 85% viability.

Hepatocytes with high-fat and low-fat content were isolated from steatotic livers^42^. Viable hepatocytes with different lipid contents were separated from dead hepatocytes and non-parenchymal cell types in the cell suspension using Percoll^®^ gradient and differential centrifugation. The hepatocytes were resuspended, washed, and assessed for cell number and viability using trypan blue and a hemacytometer and immediately plated for experiments or pelleted and stored at −80°C for further RNA or protein extractions used in RNA-Seq and proteomic/phosphoproteomic analyses.

HepG2, HLE, and Huh6 cell lines were cultured using DMEM with 10% heat-inactivated FBS and Penicillin-Streptomycin. For mouse primary hepatocytes, 100,000 cells/well in a p24 plate using attachment medium (William’s Medium with Glutamax, 10% Foetal Bovine Serum (FBS), 1% Penicillin-Streptomycin, and 10 mM HEPES). After attachment, the media was replaced with a maintenance medium (William’s E Medium with Glutamax, 10% FBS, 1% Penicillin-Streptomycin, 1% non-essential amino acids, 10mM HEPES, and 5µM hydrocortisone). Cell death was measured using SYTOX green (Thermo Fisher Scientific).

### LC-MS analysis of total proteome and phosphoproteome changes

The cell pellets originating from primary hepatocytes were lysed in cold HEN Buffer supplemented with PhosSTOP (Roche) and cOmplete™, EDTA-free Protease Inhibitor Cocktail (Roche). Lysates were precipitated twice using methanol/chloroform precipitation (Sample:Methanol:Chloroform, 4:4:1). Pellets were resuspended using 2% SDS in 50mM HEPES, and protein concentration was performed using DC protein assay (Bio-Rad Laboratories). An equal amount of protein was subjected to reduction and alkylation by incubating the samples with 5mM DTT for 1h at 37°C, followed by incubation with 20mM iodoacetamide at room temperature for 30min. Pellets were dissolved using 6M Guanidine-HCl in digestion buffer (50mM ammonium bicarbonate, 1mM CaCl2). 250µg of proteins were diluted to 0.3M Guanidine-HCl in digestion buffer and digested using trypsin (protein:trypsin, 20:1, w/w). Peptides were desalted on Supel™-Select HLB SPE Tube (Sigma-Aldrich). Eluates were evaporated under a vacuum until dryness. Peptides were dissolved in 80% acetonitrile and 0.1% TFA. Before enrichment, digestion quality control was performed using an Ultimate 3000 Nano Ultra High-Pressure Chromatography system with a PepSwift Monolithic® Trap 200µm*5mm (Thermo Fisher Scientific). One part of the sample was kept aside and dried again for total proteomic analysis. The phosphorylated peptides enrichment was performed using Fe(III)-NTAcartridges (Agilent Technologies) using the AssayMAP Bravo Platform (Agilent Technologies)^43^. Cartridges were primed and equilibrated with 0.1% TFA in ACN and 0.1% TFA, 80% ACN (loading buffer) solutions, respectively. Cartridges were then washed with loading buffer and eluted using 1% NH4OH. Peptides were immediately acidified using 10% formic acid (FA) and dried in vacuum. Both total proteome and phosphoproteome were analysed by high-resolution LC-MS/MS using an Ultimate 3000 Nano Ultra High-Pressure Chromatography system (Thermo Fisher Scientific) coupled with an Orbitrap Eclipse™ Tribrid™ Mass Spectrometer via an EASY-spray (Thermo Fisher Scientific). For the total proteome analysis, peptide separation was carried out with an Acclaim™ PepMap™ 100 C18 column (Thermo Fisher Scientific) using a 155min linear gradient from 3 to 35% of B (84% ACN, 0.1% FA) at a flow rate of 250nL/min. The peptide separation for the phosphoproteome analysis was carried out with an Acclaim™ PepMap™ 100 C18 column (Thermo Fisher Scientific) using a 155min non-linear gradient from 3 to 35% of B (0 min, 3% B; 135min, 30% B; 155min, 42% B; B:84% ACN, 0.1% FA) at a flow rate of 250nL/min. Both were analysed using the Orbitrap Eclipse™ operated in a DDA mode. MS1 survey scans were acquired from 300 to 1,500m/z at a resolution 120,000 using the Orbitrap mode. MS2 scans were carried with high-energy collision-induced dissociation (HCD) at 32% using the Normal speed IonTrap mode. Data were evaluated with Proteome Discoverer software using 10ppm for precursor mass tolerance, 0.5Da for the fragment mass tolerance, specific tryptic digest, and a maximum of 3 missed cleavages. Carbamidomethylation (+57.021464Da) on C was added as a fixed modification. N-term Acetylation (+42.010565Da) and methionine oxidation (+15.994915Da) were added as variable modifications. Phosphorylation (+79.966331) on S, T, and Y was added as variable modification only for the phospho-proteome analysis in addition to other mentioned modifications. Peptide-spectrum matches and proteins were filtered at FDR 1%. Protein abundancies (total proteome) were normalized using TIC. Phosphorylated peptide abundancies were normalized using eigenMS^44^ with R studio.

### Adenoviral infection

Adenoviral-mediated hepatic PTPRK overexpression was performed by retro-orbital injection of 1.8×10^9^ PFU (Ad-*Ptprk*, contruct Ad-m-PTPRK, SKU: ADV-269821) in 200µL of PBS, and Adv-CMV-Null was used as control (Ad-control, #1300).

In primary mouse hepatocytes, we used adenoviral vectors to achieve PTPRK overexpression (Ad-*Ptprk*, contruct Ad-m-PTPRK, SKU: ADV-269821) and silencing (Ad- shRNA *Ptprk*, contruct Ad-GFP-U6-m-PTPRK-shRNA, SKU: shADV-269821). The cells were used for the experiments 48h after the exposure to the vectors. The vectors were obtained from Vector Biolabs.

### In vitro RNA interference and colony formation assay

To induce PTPRK knockdown, we transfected HepG2, HLE, and Huh6 cell lines with siRNAs targeting PTPRK or a negative control siRNA (working concentration 30nmol/L; QIAGEN). The delivery of siRNA was achieved using Lipofectamine™ RNAiMAX Transfection Reagent (Thermo Fisher Scientific) in Opti-MEM™ I Reduced Serum Medium (Thermo Fisher Scientific). The siRNA target sequences are detailed in **Supplementary Table 2**.

48-h after siRNA transfection, the cells underwent trypsinization to attain a single-cell suspension. For the colony formation assays, 2,000 cells were seeded into P6 plates. After 1- 2 weeks, depending on the specific cell line, the resultant colonies were fixed with 4% PFA solution and staining with 0.5% crystal violet.

### Dephosphorylation assay

The dephosphorylation assay employed pTyr-enriched lysates obtained from primary mouse hepatocytes treated with pervanadate^13^. The hepatocytes were incubated with a recombinant PTPRK intracellular domain (PTPRK-ICD, 150nM final concentration). The reaction was stopped after 90-min using SDS, and the resulting samples were subjected to immunoblot analysis of phospho-FBP1 (pY265).

### Extracellular acidification rates measurement during glycolytic stress test

Glycolytic rates were evaluated using the XFp Flux Analyzer from Seahorse Bioscience (Agilent Technologies). Primary hepatocytes were plated in the Seahorse plates (10,000 cells/well). After attachment and adenoviral treatments, the cells were allowed to equilibrate in XF Base media (glucose-free, Seahorse Bioscience) supplemented with 2mM glutamine at 37°C for 1h in a CO2-depleted incubator. The XF Base media was refreshed again immediately before starting the ECAR measurements. The glycolytic stress test was conducted adding glucose (10mM), oligomycin (10μM), and 2-DG (100mM, divided into two consecutive injections of 50mM). After the test, the medium was removed, and the cells were immediately collected in 50μl of cell lysis buffer (Cell Signaling Technology) supplemented with Halt protease and phosphatase inhibitor cocktail (Thermo Fisher Scientific). The cells were stored at −80°C for posterior protein measurements and immunoblot analysis.

### Western blotting

RIPA buffer (Cell Signaling Technology) was used to extract total protein lysates from tissues, while cell total protein lysates were prepared using Cell Lysis Buffer (Cell Signaling Technology). Both lysis buffers were supplemented with Halt protease and phosphatase inhibitor cocktail (Thermo Fisher Scientific). Protein quantification was performed using a BCA protein assay kit (Thermo Fisher Scientific). Separated by polyacrylamide gels, 20–50µg of protein lysate was subsequently transferred to a 0.22µM nitrocellulose membrane (Bio-Rad Laboratories). Primary antibodies (**Supplementary Table 3**) were diluted in milk-blocking buffer. Detection of proteins employed goat anti-rabbit IgG (Dako Agilent), goat anti-mouse IgG (Dako Agilent), and Peroxidase AffiniPure Donkey Anti-Human IgG (Jackson ImmunoResearch) secondary antibodies. Immunoreactive bands were detected using a Western blot imaging system (Amersham ImageQuant 800 Western blot imaging system, Cytiva Life Science).

### RNA extraction, qPCR, and transcriptomics analysis

Poly(A)+ mRNA extraction was performed with Dynabeads™ mRNA DIRECT™ Purification Kit (Thermo Fisher Scientific). Reverse transcription was carried out with a reverse transcriptase kit (Eurogentec). Quantitative real-time PCR was performed using a Bio-Rad CFX (Bio-Rad Laboratories) and SYBR Green reagents (Bio-Rad Laboratories). Probe and primer details can be found in **Supplementary Table 4.** For tissues or isolated hepatocytes with high-fat and low-fat content the total RNA was obtained using a RNeasy Mini Kit (QIAGEN) following the manufacturer’s instructions. cDNA synthesis and qPCR were performed as described above. For the transcriptomics experiments, total RNA quality analysis, library preparation, and sequencing were performed by the BRIGHTcore facility (Brussels, Belgium). Sequencing was performed on an Illumina NovaSeq 600. An average of 25 million paired-end reads of 100 nucleotides were obtained per sample. The list of up-/downregulated genes/transcripts and association with canonical pathways were determined with the use of the online Degust software with Limma/Voom and packages Bioconductor EGSEA and ComplexHeatmap in RStudio.

### Bioinformatic analysis

For the comparative analysis among healthy, steatosis and NASH conditions, the Kruskal-Wallis test was employed. The comparison analysis was conducted on mRNA expression data from the publicly available E-MEXP-3291 study, and the analysis was executed using R (version 4.2.2). The correlation analysis focused on investigating the relationship between Ppar mRNA (x-axis) and RPTPs mRNA (y-axis) using the Pearson correlation method. The data was obtained from the publicly available E-GEOD-48452 study. The analysis was performed in R (version R 4.2.2). To assess the statistical significance of the correlation, a significance level (*p* < 0.05) was set, and p-values were calculated. To study the expression of RPTPs in human liver, single-cell RNA-Seq dataset of human healthy-obese livers was obtained from^18^. The dataset along with cell annotations were downloaded from Gene Expression Omnibus (GEO), accession number GSE192740. Using the information provided by the authors, the UMAP and gene expression were plotted using Seurat^45^. Publicly available transcriptomic data (RNA-seq) corresponding to GSE164760 was downloaded from the NCBI Sequence Read Archive (SRA) in fastq format using version 3.0.0 of the *SRA Toolkit*. Adapter sequences were removed using *TrimGalore* version 0.6.0 with *Cutadapt* version 1.18^46^. The clean reads were aligned to the reference genome using the splice-aware aligner *STAR* version 020201^47^ based on the hg38 genome version. The aligned reads were quantified using *HTseq* version 0.11.0. The trimmed mean of M values method was used with *EdgeR*version 3.28.1^48^, R software (version 3.6.3).

Heatmap visualization was carried out with *ComplexHeatmap* R package and all samples were referred to the mean of the control groups, log2 transformed with trimmed standard deviation in proteomic and transcriptomic databases.

### Measurement of fructose 1,6-bisphosphate in cells using HYlight

The HYlight biosensor^22^ responds to changes in fructose 1,6-bisphosphate levels. Cells were transfected with the pCS2+_HYlight plasmid^22^ using Lipofectamine 3000 (ThermoFisher). 1h before imaging, cells were subjected to glucose starvation using XF assay medium. Live-cell imaging was conducted on a Nikon AX confocal system and a 20X objective (NA 0.8, Plan Apo λD 20x OFN 25 DIC N2) with the perfect focus system (PFS). The transfected cells were excited at 488nm and 405nm, and emission was captured using a 525/25nm emission filter. Imaging was performed at 37°C. Image processing was performed with NIS-Elements software (Nikon), single-cell regions of interest (ROIs) were manually selected. The excitation ratio F488/405 was measured for each ROI over time.

### PTP activity assay

Recombinant PTPRK intracellular domain (PTPRK ICD) was purified^13^, and the pNPP phosphatase activity assay was conducted as previously described^49^. PTP was buffer exchanged to the activity assay buffer (20mM HEPES, 100mM NaCl, pH 7.4) using an Amicron 10kDa MWCO. The assay buffer was degassed by flushing with Argon. PTP was subjected to the indicated treatments (inhibitors or vehicle) and loaded into a 96-well plate. The reaction was initiated by the addition of 15mM or the indicated concentrations of pNPP to the reaction mixture. To determine the IC50 value, the reactions were prepared by adding the inhibitor or DMSO to a reaction mixture containing PTP, incubating at room temperature for 5 min, and centrifuged at 14,000 rpm. The supernatant was transferred to a 96-well plate. The reaction was initiated by the addition of 30mM pNPP to the reaction mixture. The formation of p-nitrophenol was measured at absorbance of 405nm at 27°C on an ID5 spectrometer. Initial velocity (V0) was determined using linear regression in GraphPad Prism.

### Stem cell differentiation into HLCs

The differentiation of CRISPR/Cas12-edited hESC H1 (WiCell) into HLCs followed the protocol previously described^50^. Laminin-coated plates were prepared and stem cells were detached, seeded into the laminin-coated plates, and allowed to reach optimal confluency before differentiation. Albumin was measured in the cell culture medium by Human ALB/Serum albumin ELISA Kit (Sigma-Aldrich) and in the cell lysate by qPCR. The differentiated cells were used for glycolytic stress tests.

### Statistical analysis

The results are presented as the mean ± standard error of the mean (SEM). Student’s t-test was used for comparisons between two groups. Differences among groups were assessed by two-way ANOVA or repeated-measures ANOVA. Statistical analyses were assessed using Prism software (GraphPad Software, Inc, La Jolla, CA, USA). Sample size was predetermined based on the variability observed in prior experiments and on preliminary data. Differences were regarded as statistically significant if \**p*< 0.05; \*\**p*< 0.01; \*\*\**p*< 0.001.

### Data and Resource Availability

The RNA-Seq dataset generated during the sequencing procedure is deposited in the Gene Expression Omnibus database (access number GSE247670), the mass spectrometry proteomics and peptidomics datasets have been deposited to the ProteomeXchange Consortium via the PRIDE partner repository (access numbers PXD046949, PXD46506) and available from the corresponding author upon reasonable request.

#### Financial support

This work was supported by a European Research Council (ERC) Consolidator grant METAPTPs (Grant Agreement No. GA817940), FNRS-WELBIO grant (35112672), FNRS-PDR grant (40007740), FNRS-TELEVIE grant (40007402), and ULB Foundation. The Ministry of Science and Innovation, through its State Plan for Scientific, Technical and Innovation Research (Project PID2021-125188OB-C32) and the Generalitat Valenciana (PROMETEO/2021/059) supported the work in the Encinar laboratory. MRF is funded by European Research Council (ERC) under the European Union’s Horizon 2020 research and innovation programme (Grant Agreement No. 864921). JM is supported by a VIB grant. ENG is a Research Associate of the FNRS, Belgium.

## Acknowledgements

We thank André Dias, Madalina Popa, Erick Arroba, Mariana Nunes, Francisco Costa, Anne Van Praet, Anaïs Schaschkow (Université libre de Bruxelles) for experimental and technical support, Nicolas Baeyens (Université libre de Bruxelles) for live-imaging microscopy advice, John Koberstein (Howard Hughes Medical Institute) for the HYlight biosensor, Carlos Martinez-Cáceres (IMIB) for pathological analysis of the samples, the Consortium des Équipements de Calcul Intensif (CÉCI) for the computer cluster NIC5, and Hercules2 for providing computing facilities. We thank Cédric Blanpain (Université libre de Bruxelles), Sarah-Maria Fendt (KULeuven) and Latifa Bakiri (Medical University of Vienna) for critical reading of the manuscript.

