## Supplemental Figures and Tables for "Protein tyrosine phosphatase receptor kappa regulates glycolysis and *de novo* lipogenesis to promote hepatocyte metabolic reprogramming in obesity"

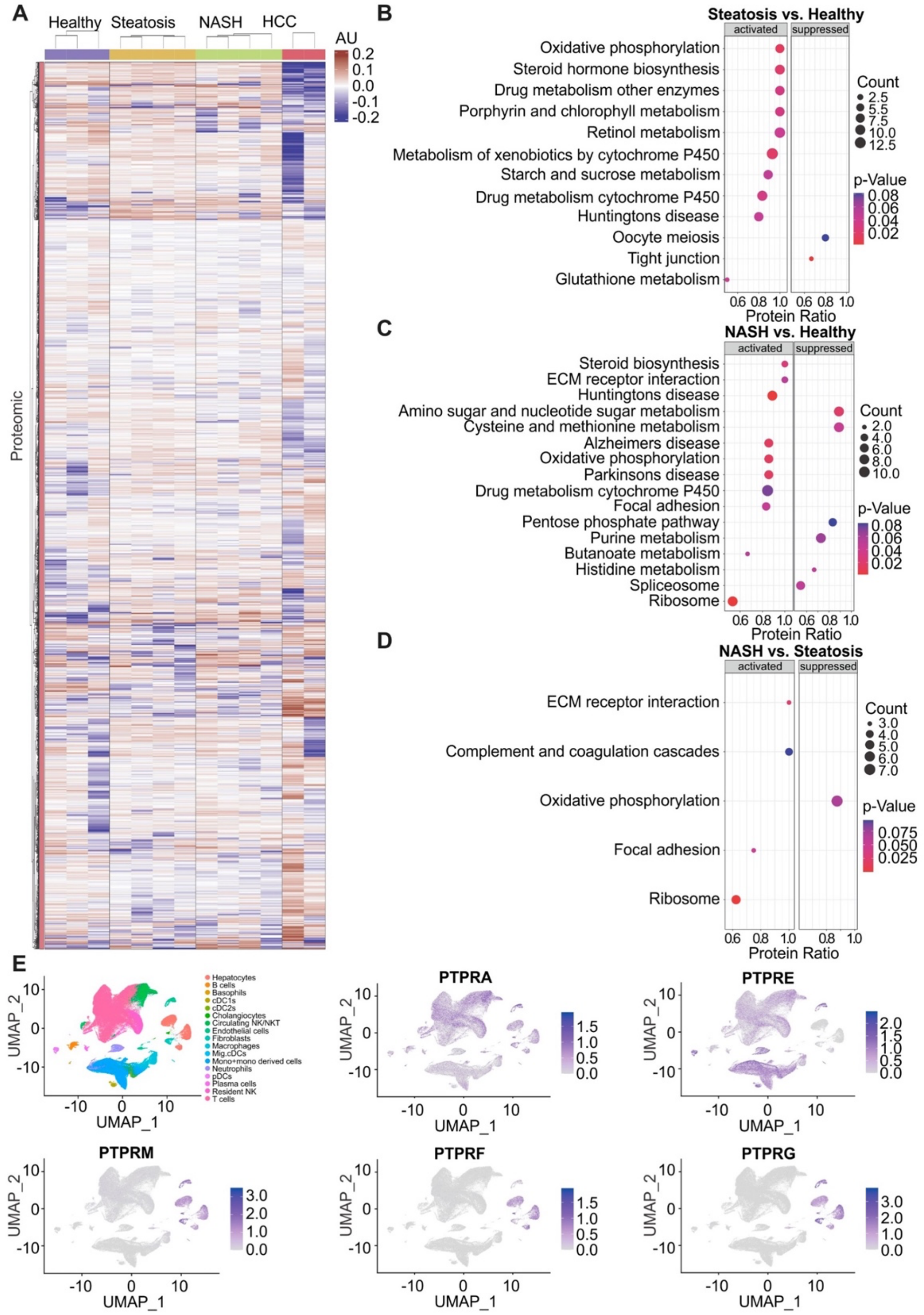


**Supplementary Figure 1: Comprehensive liver proteomics, pathway enrichment, and receptor PTPs expression in different hepatic cells.**

(**A**) Heat map displaying the hepatic proteome profile. (**B**) Proteomic KEGG pathway enrichment analysis comparing steatosis and healthy livers. (**C**) Proteomic KEGG pathway enrichment analysis comparing NASH and healthy livers. (**D**) Proteomic KEGG pathway enrichment analysis comparing NASH and steatosis. (**E**) Data extracted from GSE192740 showing UMAPs with distinct cell clusters identified in the liver and the distribution of various receptor PTPs across parenchymal and non-parenchymal hepatic cells.

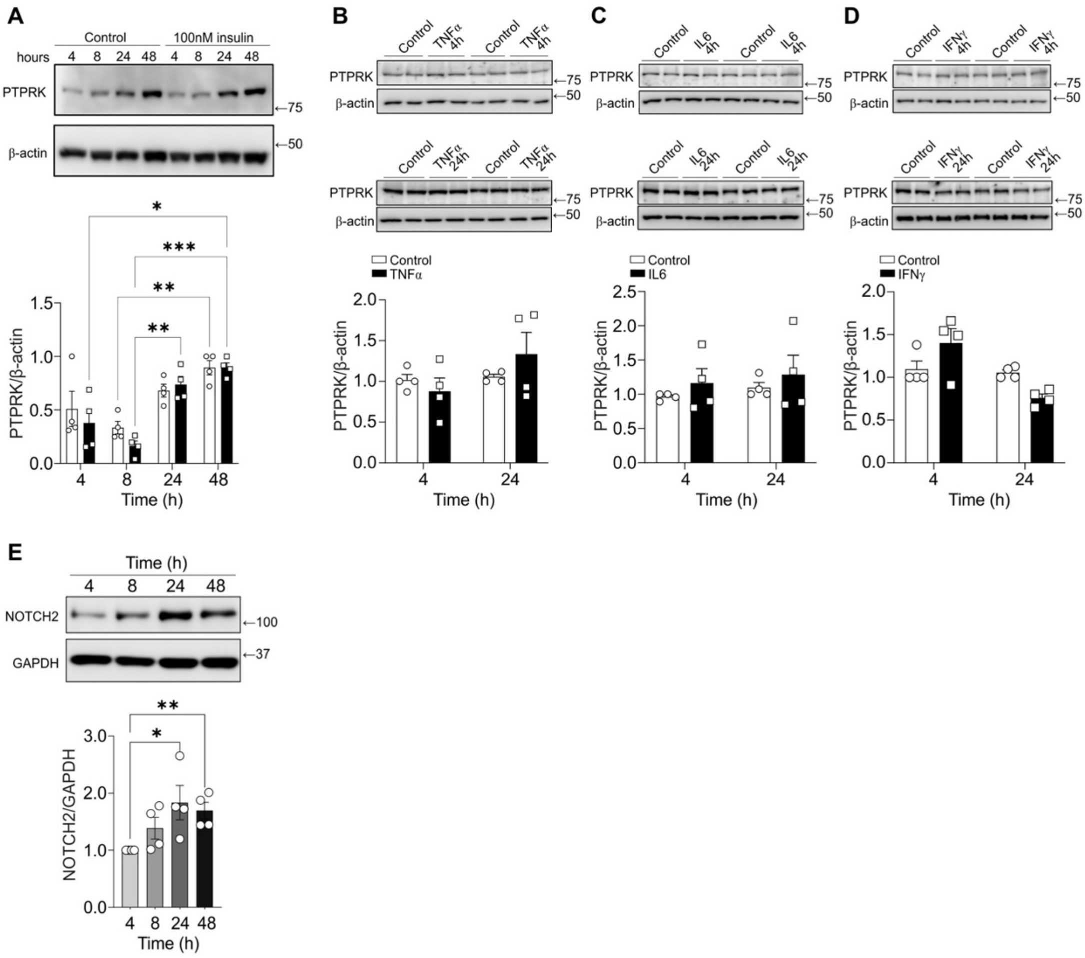


**Supplementary Figure 2: Insulin or cytokine treatment does not affect PTPRK expression and culture of primary hepatocytes enhances Notch2 expression.**

(**A**) Immunoblot analysis of PTPRK expression in mouse primary hepatocytes cultured over time treated with insulin as indicated. (**B**) Immunoblot analyses of PTPRK expression in mouse primary hepatocytes with acute (4h) and chronic (24h) TNFα treatment. (**C**) Immunoblot analyses of PTPRK expression in mouse primary hepatocytes with acute (4h) and chronic (24h) IL6 treatment. (**D**) Immunoblot analyses of PTPRK expression in mouse primary hepatocytes with acute (4h) and chronic (24h) IFNγ treatment. (**E**) Immunoblot analyses of NOTCH2 expression in mouse primary hepatocytes over time in culture. The presented data represents the average of different independent experiments and are expressed as mean±SEM. Statistical significance is indicated as ***p*<0.05, ***p*<0.01, ****p*<0.001.

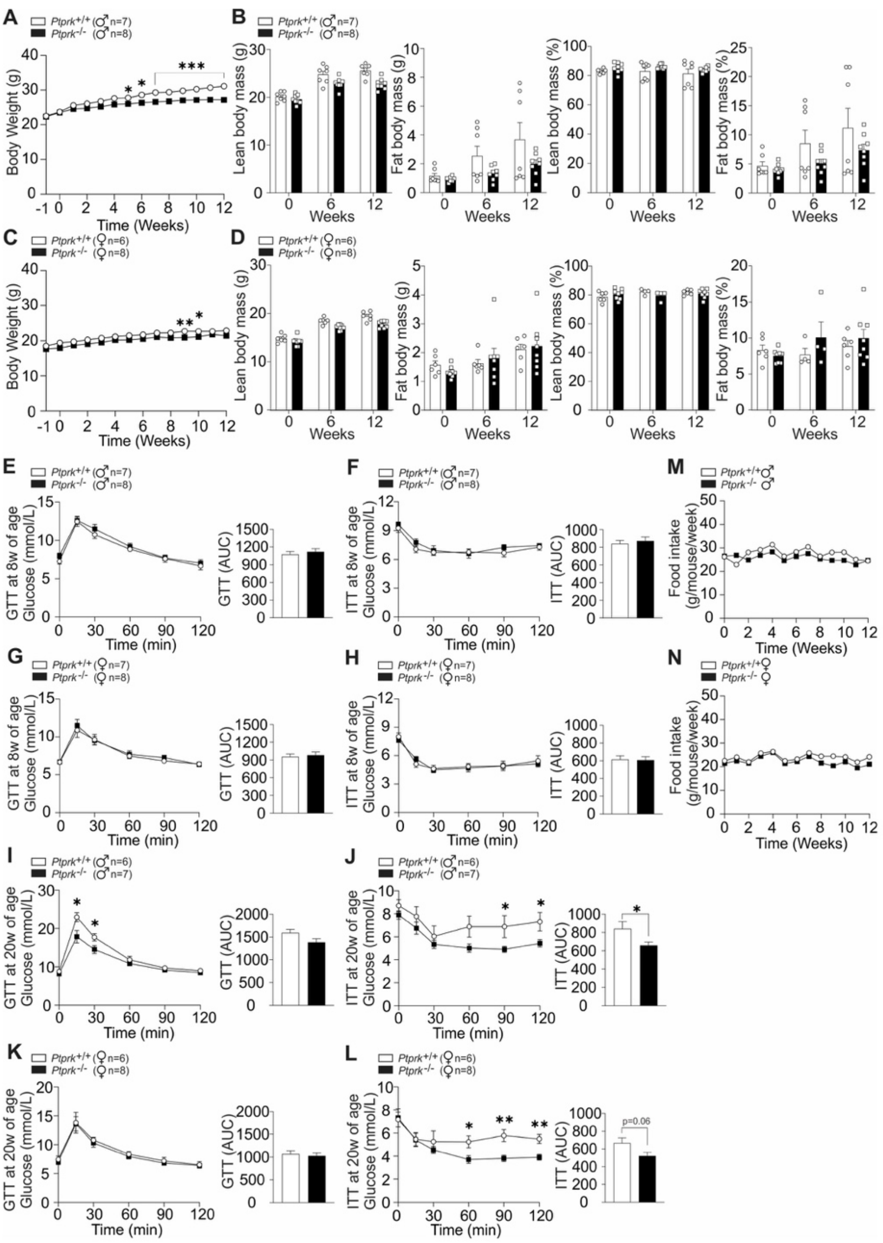


**Supplementary Figure 3: Metabolic phenotype shows minor differences in body weight and composition in *Ptprk*$^{-/-}$ and *Ptprk*$^{+/+}$ mice fed a chow diet.**

(**A**) Male (♂) *Ptprk*$^{-/-}$ and *Ptprk*$^{+/+}$ C57BL6N mice aged eight weeks, continuously fed a chow diet for additional 12 weeks, and body weight was measured weekly. (**B**) Body composition in male mice. (**C**) Female (♀) *Ptprk*$^{+/+}$ and *Ptprk*$^{-/-}$ C57BL6N mice aged eight weeks, continuously fed a chow diet for 12 weeks, and body weight was measured weekly. (**D**) Body composition of female mice. (**E**) Glucose tolerance test in male mice at 8 weeks of age. (**F**) Insulin tolerance test in male mice at 8 weeks of age. (**G**) Glucose tolerance test in female mice at 8 weeks of age. (**H**) Insulin tolerance test in female mice at 8 weeks of age. (**I**) Glucose tolerance test in male mice at 20 weeks of age. (**J**) Insulin tolerance test in male mice at 20 weeks of age. (**K**) Glucose tolerance test in female mice at 20 weeks of age. (**L**) Insulin tolerance test in female mice at 20 weeks of age. (**M**) Food intake in male mice. (**N**) Food intake in female mice. The presented data represent the average of independent experiments and are expressed as mean±SEM. Statistical significance is indicated as *$p$<0.05, **$p$<0.01, ***$p$<0.001.

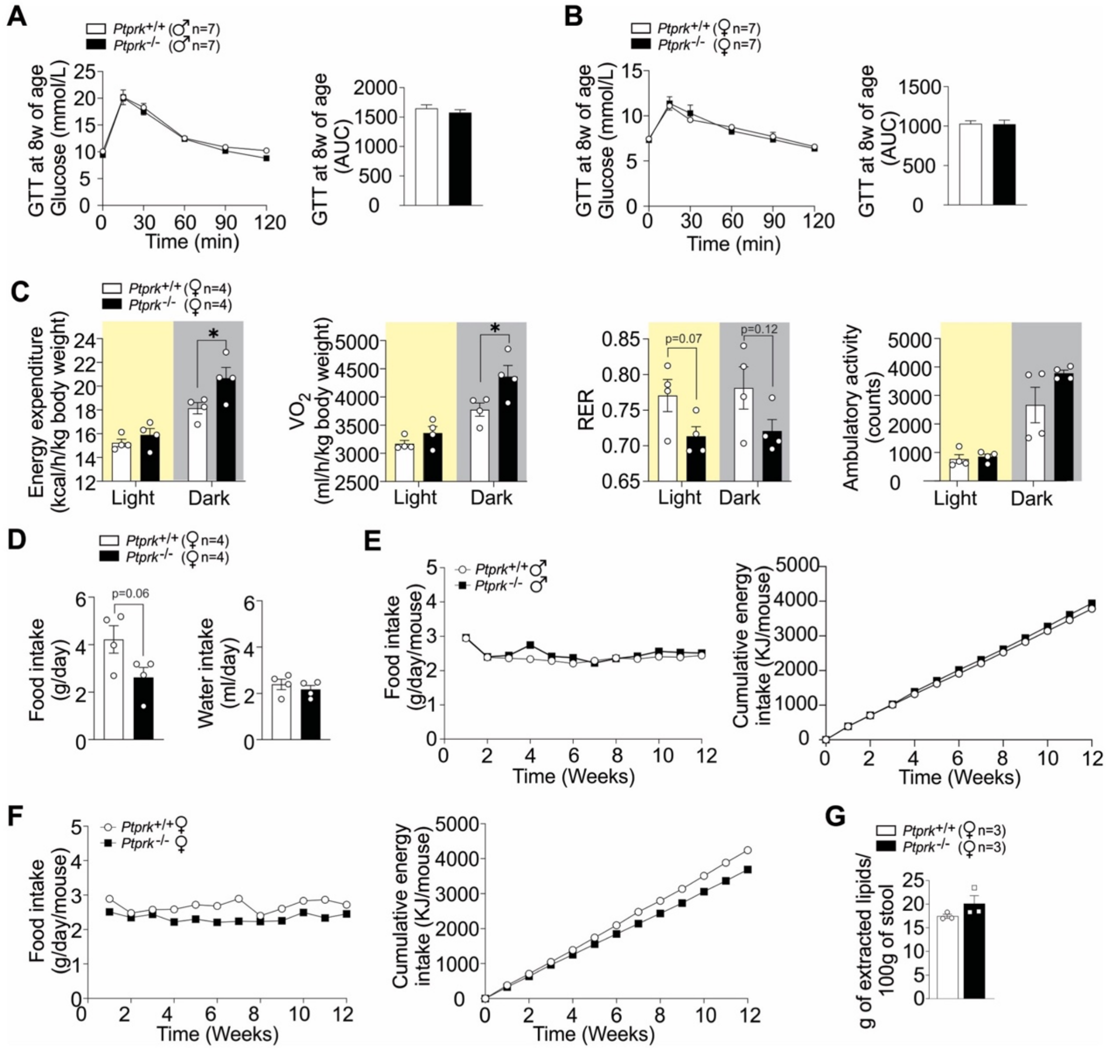


**Supplementary Figure 4: Metabolic assessments in *Ptprk*$^{-/-}$ and *Ptprk*$^{+/+}$ mice fed a high-fat, high-fructose, high-cholesterol diet.**

(**A, B**) Glucose tolerance test performed in males at 8 weeks of age (before the HFHFHCD feeding). (**C**) Energy expenditure, oxygen consumption ($VO_2$), respiratory exchange ratio ($RER=VCO_2/VO_2$) and ambulatory activity measurements in female *Ptprk*$^{+/+}$ and *Ptprk*$^{-/-}$ mice after high-fat, high-fructose, high-cholesterol diet (HFHFHCD) feeding for 8 weeks. (**D**) Daily food and water intake. (**E**) Average food intake and cumulative energy intake analyses in males HFHFHCD. (**F**) Average food intake and cumulative energy intake analyses in females HFHFCD. (**G**) Lipid extraction from stool samples collected over three consecutive days before the end of the 12-week HFHFHCD feeding experimental period. The presented data represent the average of independent experiments and is expressed as mean±SEM. Statistical significance is indicated as *$p<0.05$.

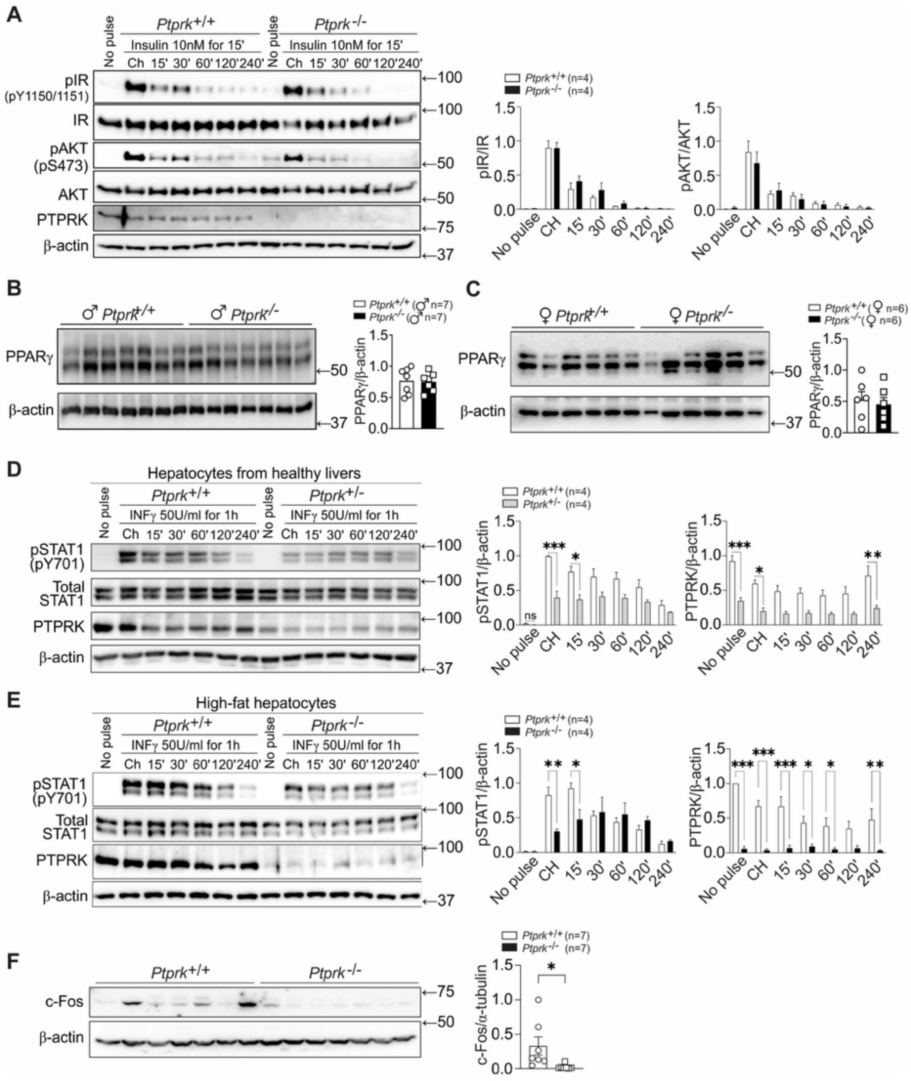


**Supplementary Figure 5: Effect of PTPRK on insulin signalling, IFNγ-induced STAT1 activation and hepatic c-Fos expression.**

(**A**) Immunoblot analysis of hepatocytes isolated from *Ptprk*$^{-/-}$, *Ptprk*$^{+/-}$, and *Ptprk*$^{+/+}$ mice cultured overnight and subjected to pulse and chase after insulin treatment. The analysis reveals that PTPRK deletion does not significantly affect hepatocyte insulin signalling, as demonstrated by pIR, IR, pAKT, AKT, and PTPRK levels in *Ptprk*$^{-/-}$ and *Ptprk*$^{+/+}$ hepatocytes isolated from healthy male mice at 8 weeks of age following a 15-min exposure to 10nM of insulin, chase (Ch). (**B, C**) Visceral epididymal (males, **B**) and uterine (females, **C**) white adipose tissues were collected for immunoblot analysis of PPARγ. (**D**) Immunoblot analysis showing lower PTPRK levels associated with decreased pSTAT1 activation in response to IFNγ in *Ptprk*$^{+/-}$ and *Ptprk*$^{+/+}$ hepatocytes after 1h exposure to 50 U/ml IFNγ. (**E**) Immunoblot analysis of IFNγ-induced STAT1 activation in *Ptprk*$^{-/-}$ and *Ptprk*$^{+/+}$ hepatocytes with high-fat content isolated from steatotic livers of male mice at 14 weeks of age following a 1-hour exposure to 50 U/ml IFNγ. (**F**) c-Fos expression was assessed by immunoblot analysis in livers from female *Ptprk*$^{-/-}$ and *Ptprk*$^{+/+}$ mice fed a high-fat, high-fructose, high-cholesterol diet for 12 weeks. The presented data represent the average of independent experiments and are expressed as mean±SEM. Statistical significance is indicated as *$p$<0.05, **$p$<0.01, ***$p$<0.001.

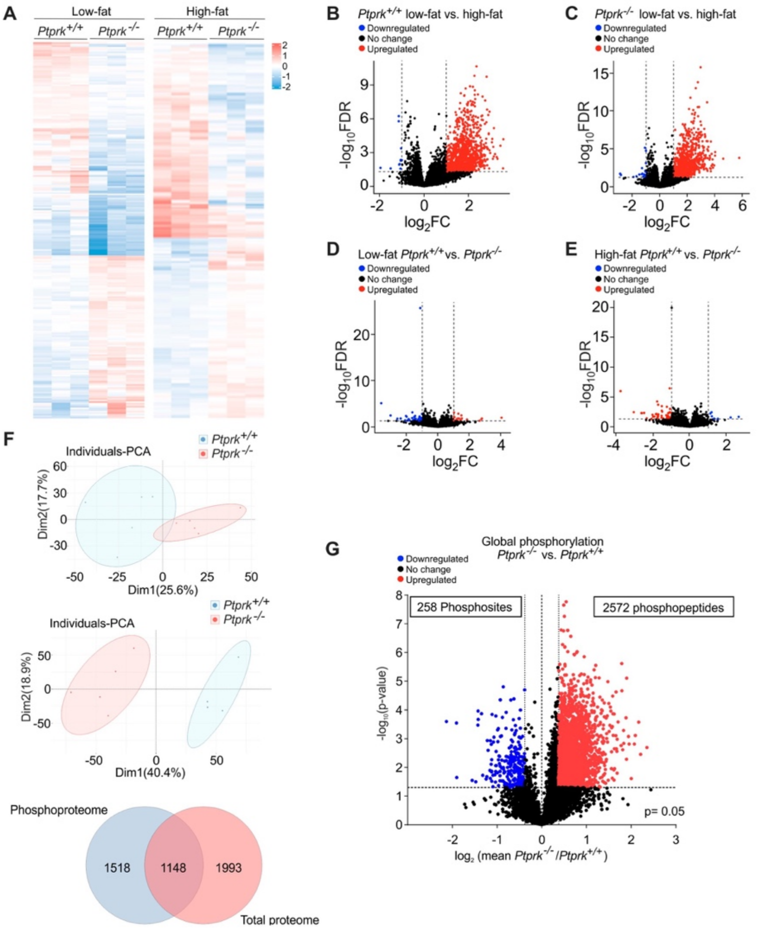


**Supplementary Figure 6: Transcriptome, proteome, and protein phosphorylation assays in primary mouse hepatocytes isolated from *Ptprk*$^{-/-}$ and *Ptprk*$^{+/+}$ mice.**

(**A**) RNA-Seq heatmap showing alterations in mRNA expression in hepatocytes with low- and high-fat content. (**B**) Volcano plot showing the quantification of transcripts in high-fat and low-fat *Ptprk*$^{+/+}$ hepatocytes. (**C**) Volcano plot showing the quantification of transcripts in high-fat and low-fat *Ptprk*$^{-/-}$ hepatocytes. (**D**) Volcano plot displaying the quantification of transcripts between *Ptprk*$^{-/-}$ and *Ptprk*$^{+/+}$ hepatocytes with low fat content. (**E**) Volcano plot displaying the quantification of transcripts between *Ptprk*$^{-/-}$ and *Ptprk*$^{+/+}$ hepatocytes with high-fat content. (**F**) PCA analysis depicting changes in the total proteome (top part) and phosphoproteome (middle part) in *Ptprk*$^{-/-}$ and *Ptprk*$^{+/+}$ hepatocytes. Principal component analysis (PCA) examination revealed that the PCA scores for PC1 (Dim1) and PC2 (Dim2) generated a biplot that segregated into two distinct groups. PC1 and PC2 collectively explained 43.3% of the variability between *Ptprk*$^{-/-}$ and *Ptprk*$^{+/+}$ samples, with PC1 accounting for 25.6% and PC2 for 17.7% of the variance, respectively. A similar analysis for the phosphoproteomic data showed a more pronounced difference, with PCA scores for PC1 (Dim1) and PC2 (Dim2) producing a biplot that distinctly separated two groups. In this case, PC1 and PC2 together explained 59.3% of the variability between *Ptprk*$^{-/-}$ and *Ptprk*$^{+/+}$ samples, with PC1 contributing 40.4% and PC2 contributing 18.9% of the variance. One group comprised *Ptprk*$^{+/+}$ hepatocytes, while the other group consisted of *Ptprk*$^{-/-}$ hepatocytes, both showing strong alignment along principal component 2 (PC2). Venn diagram illustrating proteins identified in the phosphoproteome and total proteome (bottom part). (**G**) Volcano plot showing the quantification of phosphosites between *Ptprk*$^{-/-}$ and *Ptprk*$^{+/+}$ hepatocytes with high-fat content.

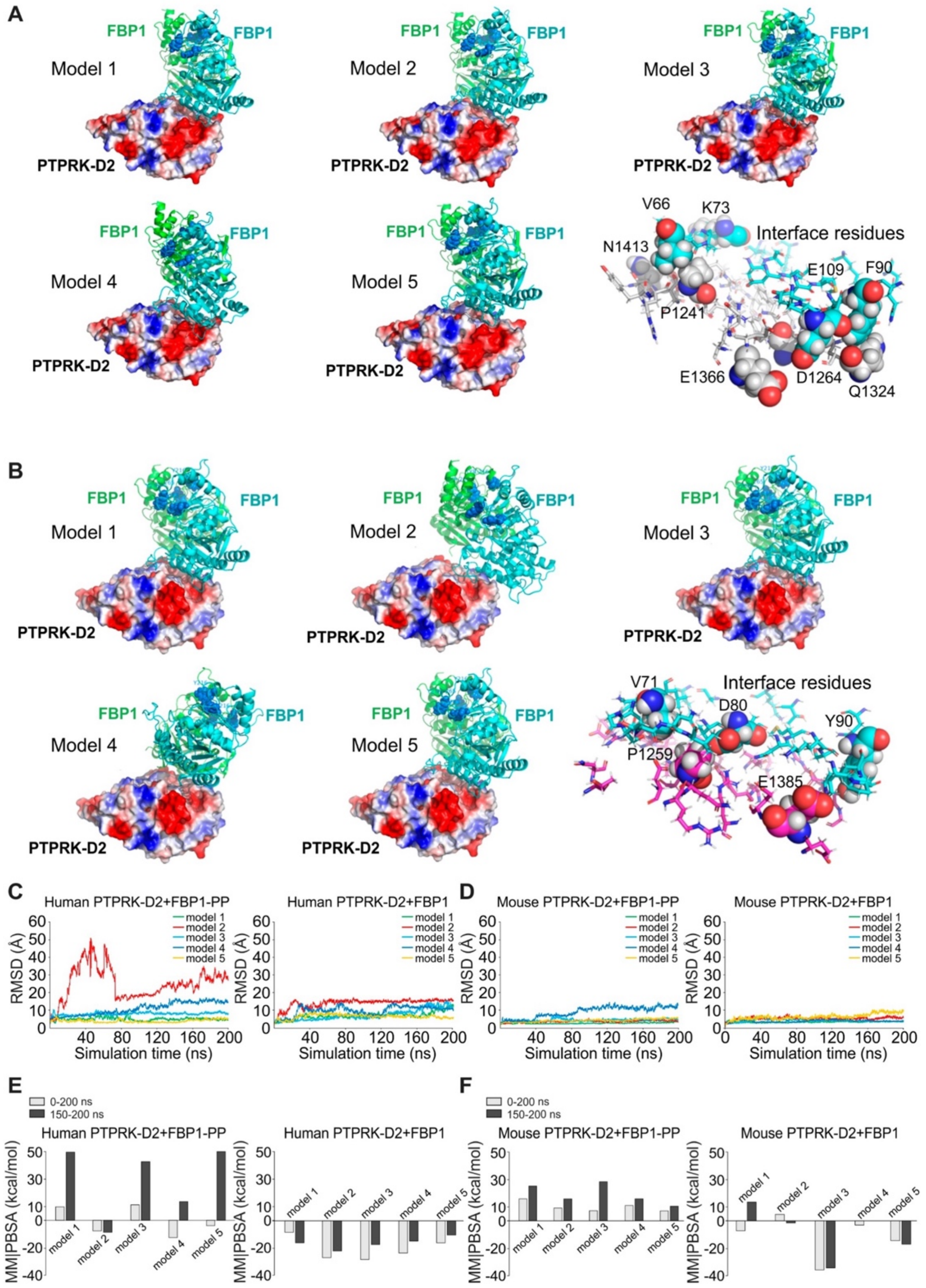


**Supplementary Figure 7: Computational modelling of the interaction between phosphorylated FBP1 dimer (PP-FBP1) and PTPRK-D2 non-catalytic domain.**
(**A-B**) The images show the surface of the human (**A**) and mouse (**B**) PTPRK-D2 non-catalytic domain and with 5 different models of the interaction between PTPRK and FBP1 (backbone of dimer are shown in green and cyan for each monomer) and the predicted interface residues participating in the PTPRK-FBP1 interaction. The colour coding represents the hydrophobicity of the amino acid side chains on the surface. (**C-D**) The RMSD values in Å for the five interaction models were calculated during the 200-ns MD simulation for human PTPRK (**C**) and mouse PTPRK (**D**). (**E-F**) The solvation binding energy (MM|PBSA, kcal/mol) values for the time intervals of 0-200 ns and the final 150-200 ns of FBP1-PTPRK D2 domain interaction simulation for phosphorylated FBP1 (PP-FBP1, pY265 and pY245) and non-phosphorylated FBP1 are presented for human (**E**) and mouse (**F**).

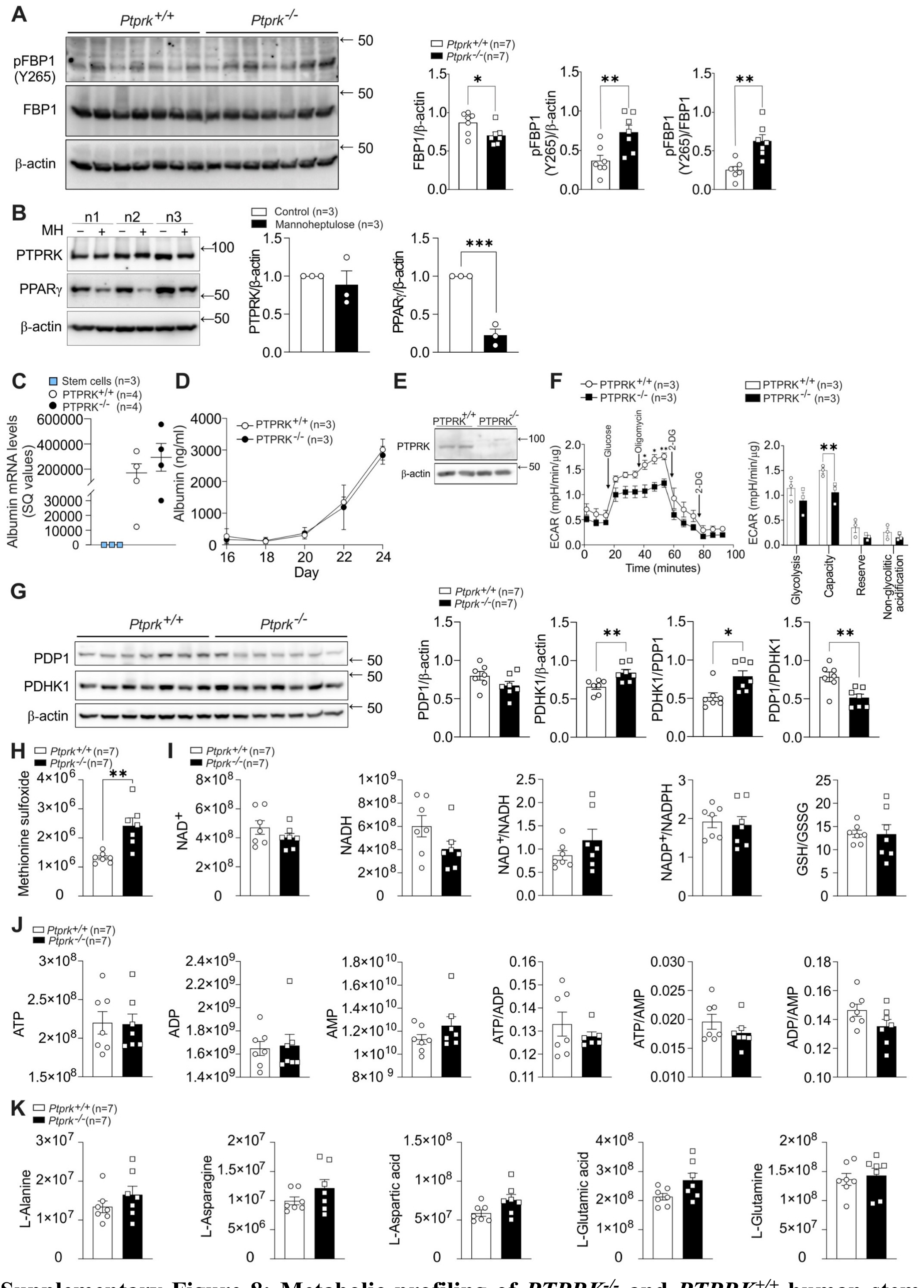


**Supplementary Figure 8: Metabolic profiling of *PTPRK*$^{-/-}$ and *PTPRK*$^{+/+}$ human stem cell-derived hepatocyte-like cells and metabolite analysis in livers from *Ptprk*$^{-/-}$ and *Ptprk*$^{+/+}$ mice.** (**A**) Liver samples from *Ptprk*$^{-/-}$ and *Ptprk*$^{+/+}$ female mice fed a 12-week high-fat, high-fructose, high-cholesterol diet (HFHFHCD) were extracted and processed for immunoblot analysis to evaluate the expression of pFBP1 (Y265) and total FBP1 levels. **(B)** Primary mouse hepatocytes were exposed to mannoheptulose (MH) as indicated and

PTPRK/PPARγ assessed by immunoblot. **(C)** Human embryonic stem cells CRISPR-Cas12a-edited to delete PTPRK were differentiated into hepatocyte-like cells. Albumin mRNA levels were measured in stem cells, *Ptprk*$^{-/-}$ and *Ptprk*$^{+/+}$ hepatocyte-like cells. (**D**) Albumin levels were quantified over the course of the differentiation process using enzyme-linked immunosorbent assay (ELISA). (**E**) Immunoblot analysis was performed after differentiation into hepatocyte-like cells. (**F**) A glycolysis stress test was conducted in *PTPRK*$^{-/-}$ and *PTPRK*$^{+/+}$ hepatocyte-like cells. Real-time measurements of the extracellular acidification rate (ECAR) in response to glycolytic modulators were recorded. (**G**) Liver samples from *Ptprk*$^{-/-}$ and *Ptprk*$^{+/+}$ female mice fed a 12-weeks HFHFHCD were extracted and processed for immunoblot analysis to evaluate the expression of PDP1 and PDHK1.(**F-K**) Livers were harvested from *Ptprk*$^{-/-}$ and *Ptprk*$^{+/+}$ female mice fed a 12-week HFHFHCD and metabolite analysis was conducted using mass spectrometry: (**H**) methionine sulfoxide, (**I**) redox status indicators $NAD^+$, NADH, $NADP^+$/NADPH and reduced glutathione (GSH) to oxidized glutathione (GSSG) ratios (**J**) energy status indicators ATP, ADP, AMP, and their ratios and (**K**) levels of amino acids. Metabolites are presented as raw abundances corrected for sample weight. The presented data represent the average of independent experiments and are expressed as mean±SEM. Statistical significance is indicated as *$p<0.05$, **$p<0.01$, ***$p<0.001$.

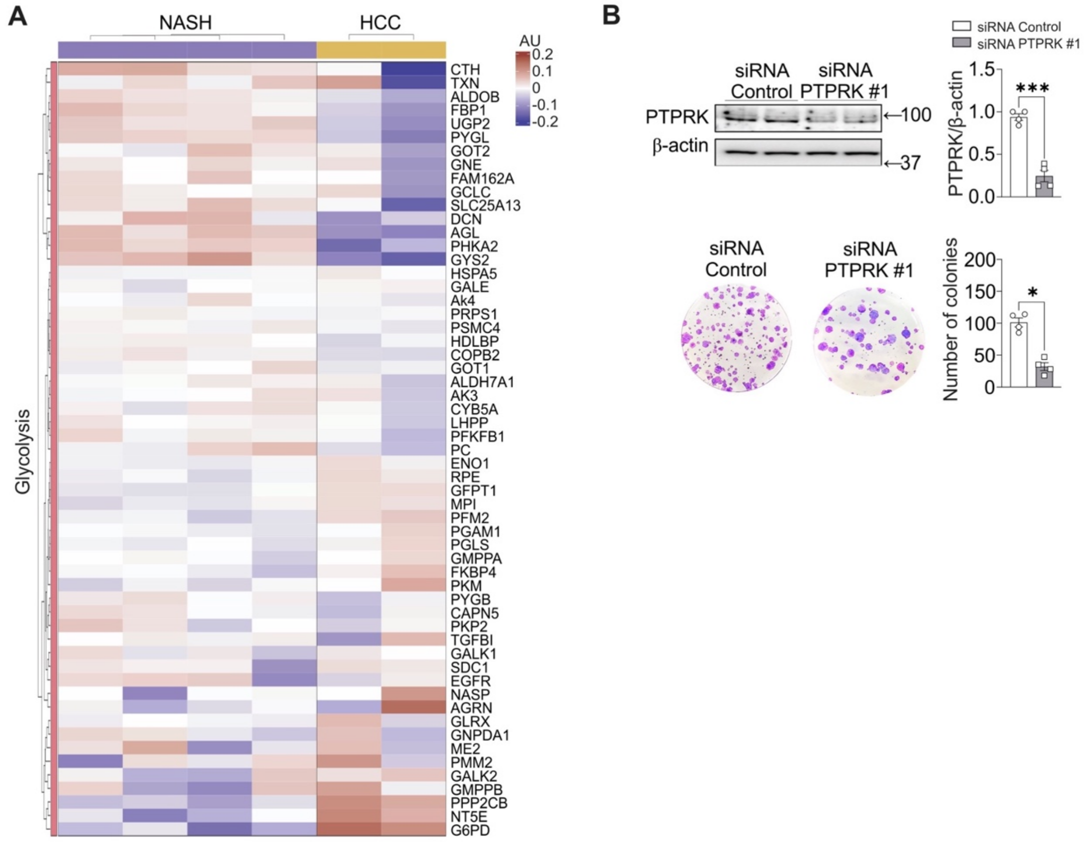


**Supplementary Figure 9: Glycolysis-related protein expression heatmaps in NASH and HCC human liver samples, and PTPRK knockdown in hepatoma cell colony formation.** (**A**) Heatmaps were generated to illustrate the expression of proteins involved in glycolysis/gluconeogenesis in human liver samples from individuals with NASH or HCC. (**B**) Huh6 cells were transfected with siRNA targeting PTPRK or siRNA control. Transfection efficiency was confirmed by immunoblot analysis, and colony-forming capacity was assessed using crystal violet staining for colony number counting. The presented data are expressed as mean±SEM, with statistical significance indicated as *$p$<0.05, ***$p$<0.001.

| ID | Biopsy health status | Age | Weight (kg) | Height (cm) | BMI (kg/m$^2$) | Type 2 diabetes | Gender | Experiment |
|---|---|---|---|---|---|---|---|---|
| CHR01 | Healthy liver | 80 | 46 | 175 | 15 | No | M | MS/IHC |
| CHR02 | Healthy liver | 66 | 62 | 150 | 28 | No | F | MS/IHC |
| CHR03 | Healthy liver | 28 | 66 | 170 | 23 | No | F | IHC |
| CHR04 | Healthy liver | 63 | 66 | 165 | 24 | No | F | MS/IHC |
| CHR08 | Steatosis | 53 | 86 | 185 | 25 | Yes | F | MS/IHC |
| CHR09 | Steatosis | 39 | 80 | 158 | 32 | No | F | MS/IHC |
| CHR10 | Steatosis | 56 | 67 | 173 | 22 | No | M | MS/IHC |
| CHR11 | Steatosis | 36 | 82 | 160 | 32 | No | F | MS/IHC |
| CHR12 | Steatosis | 61 | 76 | 159 | 30 | No | F | IHC |
| CHR13 | Steatosis | 68 | 115 | 168 | 41 | Yes | M | IHC |
| CHR14 | NASH/Cirrhosis | 61 | 113 | 179 | 35 | Yes | M | MS/IHC |
| CHR16 | NASH/Cirrhosis | 46 | 79 | 156 | 32 | No | F | MS/IHC |
| CHR17 | NASH/Cirrhosis | 48 | 76 | 160 | 30 | No | M | MS/IHC |
| CHR18 | NASH/Cirrhosis | 60 | 80 | 149 | 36 | Yes | F | MS/IHC |
| CHR19 | NASH | 46 | 80 | 157 | 32 | Yes | F | MS |
| CHR20 | NASH | 45 | 85 | 156 | 35 | Yes | F | MS |
| CHR05 | HCC | 83 | 79 | 175 | 26 | No | M | MS |
| CHR06 | HCC | 76 | 76 | 165 | 28 | No | M | MS |
| CHR07 | HCC | 43 | 55 | 170 | 19 | No | M | MS |

**Supplementary table 1:** Clinical and demographic characteristics of patients from whom liver biopsies were collected and used for mass spectrometry (MS) or PTPRK immunohistochemistry (IHC) analysis shown in Figure 1.

| siRNA | Source or reference | Identifiers | Sense (5´->3´) | Antisense (5´->3´) |
|---|---|---|---|---|
| siRNA PTPRK #1 | Ambion by Life Technologies | s11559 | CCAGUAGCCCAG ACUAAGAtt | UCUUAGUCUGGG CUACUGGta |
| siRNA PTPRK #2 | Ambion by Life Technologies | s11558 | CAGCUAUAGCAG UAUAAGAtt | UCUUAUACUGCUA UAGCUGat |
| siRNA Control | Quiagen | 1027281 | - | - |

**Supplementary table 2:** List of siRNAs used for RNA interference in the present study.

| Antibody | Designation | Source or reference | Identifiers | Additional information |
|---|---|---|---|---|
| β-actin | Mouse monoclonal anti-beta-actin | Sigma-Aldrich | Cat# A1978 | Western blot 1:5000 |
| GAPDH | Rabbit polyclonal anti-glyceraldehyde-3-phoshphate dehydrogenase | R&D Systems | Cat# 2275-PC-100 | Western blot 1:5000 |
| α-tubulin | Mouse monoclonal anti-alpha-tubulin | Sigma-Aldrich | Cat# T5168 | Western blot 1:5000 |
| Vinculin | Rabbit polyclonal anti-vinculin | Cell Signaling Technology | Cat# 4650 | Western blot 1:1000 |
| pY | Mouse monoclonal anti-phospho-tyrosine antibody | Cell Signaling Technology | Cat# 9411 | IP 1:50<br>Western blot 1:1000 |
| PTPRK | Human monoclonal anti-PTPRK | Fearnley et al. 2019, Kindly provided by Hayley Sharpe | 2 .H4 | Western blot 1:1000 |
| PTPRK | Rabbit polyclonal anti-PTPRK | Thermo Fisher Scientific | Cat# PA5-104089 | IHC 1:100 |
| PPARγ | Rabbit monoclonal anti- PPARγ | Cell Signaling Technology | Cat# 2443 | Western blot 1:500 |
| pIR | Rabbit monoclonal anti-phospho-IGF-I Receptor β (Tyr1135/1136)/Insulin Receptor β (Tyr1150/1151) | Cell Signaling Technology | Cat# 3024S | Western blot 1:500 |
| IR | Rabbit monoclonal anti-Insulin Receptor β | Cell Signaling Technology | Cat# 3025 | Western blot 1:1000 |
| pAKT | Rabbit monoclonal anti-phospho-AKT (Ser473) | Cell Signaling Technology | Cat# 4060 | Western blot 1:1000 |
| AKT | Mouse monoclonal anti-AKT (pan) | Cell Signaling Technology | Cat# 2920S | Western blot 1:1000 |
| ACC | Mouse monoclonal anti-acetyl CoA carboxylase | Cell Signaling Technology | Cat# 3676S | Western blot 1:1000 |
| FASN | Rabbit monoclonal anti-fatty acid synthase | Cell Signaling Technology | Cat# 3180S | Western blot 1:1000 |
| SREBP1 | Mouse monoclonal anti-sterol regulatory element binding protein 1 | Santa Cruz Biotechnology | Cat# sc-13551 | Western blot 1:1000 |
| ChREBP1α | Rabbit polyclonal anti-Carbohydrate-responsive element-binding protein alpha | Cell Signaling Technology | Cat# 58069 | Western blot 1:1000 |
| pFBP1 (Y265) | Rabbit polyclonal anti-Phospho-fructose 1,6 bisphosphatase 1 (Tyr265) | Thermo Fisher Scientific | Cat# PA5-105335 | Western blot 1:1000 |
| FBP1 | Rabbit polyclonal anti-fructose 1,6 bisphosphatase 1 | Cell Signaling Technology | Cat# 59172 | Western blot 1:1000 |
| PDP1 | Rabbit monoclonal anti-pyruvate dehydrogenase phosphatase 1 | Cell Signaling Technology | Cat# 65575S | Western blot 1:1000 |
| PDHK1 | Rabbit monoclonal anti-pyruvate dehydrogenase kinase 1 | Cell Signaling Technology | Cat# 3820S | Western blot 1:1000 |
| c-Fos | Rabbit polyclonal anti-protein c-Fos | Cell Signaling Technology | Cat# 4384S | Western blot 1:1000 |
| Notch2 | Rabbit monoclonal anti-Neurogenic locus notch homolog protein 2 | Cell Signaling Technology | Cat# 5732S | Western blot 1:1000 |
| HIF-2α | Rabbit monoclonal anti-Hypoxia-inducible factor 2 alpha | Cell Signaling Technology | Cat# 57921S | Western blot 1:500 |

**Supplementary table 3:** Antibodies used in the present study.

| Gene name | qPCR Fw | qPCR Rv | STD Fw | STD Rv |
|---|---|---|---|---|
| Tnf | AGGCACTCCCCCAAAAGATG | TGAGGGTCTGGGCCATAGAA | ATGAGCACAGAAAGCATGATC | TACAGGCTTGTCACTCGAATT |
| Pparγ1 | CCAAGAATACCAAAGTGCGATCA | AAAACCCTTGCATCCTTCACAA | GCTCCAAGAATACCAAAGTGCGA | AACCTGATGGCATTGTGAGACA |
| Pparγ2 | TGCCTATGAGCACTTCACAAG | TCTACTTTGATCGCACTTTGGTA | AGCATGGTGCCTTCGCTGAT | GCCCAAACCTGATGGCATTGTG |
| Cyp7a1 | TAAGGAGAAGGAAAGTAGGTGAAC | CCAAATGCCTTCGCAGAAGTAG | TGGAATAAGGAGAAGGAAAGTAGG | TCCAAATGCCTTCGCAGAAGTA |
| Gapdh | AGTTCAACGGCACAGTCAAG | TACTCAGCACCAGCATCACC | ATGACTCTACCCACGGCAAG | TGTGAGGGAGATGCTCAGTG |
| Fasn | CACAGTGCTCAAAGGACATGCC | CACCAGGTGTAGTGCCTTCCTC | ACTTCCTCTGGGATGTGCCT | GTCAGCACTGCTCTCGTTGA |
| Pck1 | GCAGTGAGGAAGTTCGTGGA | GTGAGAGCCAGCCAACAGT | GCTGCATAACGGTCTGGACT | ATACATGGTGCGGCCTTTCAT |
| Cd36 | TAATGGCACAGACGCAGC | GGTTGTCTGGATTCTGGAGGG | TAATGGCACAGACGCAGC | ACATCACCACTCCAATCCCA |
| Cpt1α | GCTGATGACGGCTATGGTGT | AAAGCGGTGTGAGTCTGTCT | CCACAACAACGGCAGAGCA | TCAGGAGCAACACCTATTCATTTG |
| Scd1 | TGCCGTGGGCGAGGG | ACACCCCGATAGCAATATCCA | TGATGTTCCAGAGGAGGTACTACA | AATGCATCATTAACACCCCGA |
| Ptprk | TCGTGATTGGCTTCGGG | CCAGCATTGACCTCCACA | TGAAGGAGAATGACACCCAC | TCTGAAAGAGGCAGCAAATCT |
| Acly | TTCGTCAAACAGCACTTCC | ATTTGGCTTCTTGGAGGTG | ACACCATCATCTGTGCTCGG | ATCCCAGGGGTGACGATACA |
| Acaca(Acc) | AGCCAGAAGGGACAGTAGAA | CTCAGCCAAGCGGATGTAAA | GCGCTTACATTGTGGATGGC | AAGCCTTCACTGTGCCTTCA |
| Alb(human) | GAAAAGTGGGCAGCAAATGT | GGTTCAGGACCACGGATAGA | GAGCAGCTTGGAGAGTACAA | GTTTCAGCATTAAACTCTTT |

**Supplementary table 4:** Primer sequences used for RT-PCR experiments.
